# A Composite Biomarker Signature of Type 1 Diabetes Risk Identified via Augmentation of Parallel Multi-Omics Data from a Small Cohort

**DOI:** 10.1101/2024.02.09.579673

**Authors:** Oscar Alcazar, Sung-Ting Chuang, Gang Ren, Mitsunori Ogihara, Bobbie-Jo M. Webb-Robertson, Ernesto S. Nakayasu, Peter Buchwald, Midhat H. Abdulreda

## Abstract

**Background:** Biomarkers of early pathogenesis of type 1 diabetes (T1D) are crucial to enable effective prevention measures in at-risk populations before significant damage occurs to their insulin producing beta-cell mass. We recently introduced the concept of integrated parallel multi-omics and employed a novel data augmentation approach which identified promising candidate biomarkers from a small cohort of high-risk T1D subjects. We now validate selected biomarkers to generate a potential composite signature of T1D risk

**Methods:** Twelve candidate biomarkers, which were identified in the augmented data and selected based on their fold-change relative to healthy controls and cross-reference to proteomics data previously obtained in the expansive TEDDY and DAISY cohorts, were measured in the original samples by ELISA

**Results:** All 12 biomarkers had established connections with lipid/lipoprotein metabolism, immune function, inflammation, and diabetes, but only 7 were found to be markedly changed in the high-risk subjects compared to the healthy controls: ApoC1 and PON1 were reduced while CETP, CD36, FGFR1, IGHM, PCSK9, SOD1, and VCAM1 were elevated

**Conclusions:** Results further highlight the promise of our data augmentation approach in unmasking important patterns and pathologically significant features in parallel multi-omics datasets obtained from small sample cohorts to facilitate the identification of promising candidate T1D biomarkers for downstream validation. They also support the potential utility of a composite biomarker signature of T1D risk characterized by the changes in the above markers.

## INTRODUCTION

It is well established that type 1 diabetes (T1D) results from the autoimmune attack against the insulin producing beta cells in the islets of Langerhans of the endocrine pancreas. The specific etiology of this anti-beta cell autoimmunity, however, remains elusive [1, 2]. Aside from a well-known genetic predisposition attributed largely to certain human leukocyte antigen (HLA) genes in the major histocompatibility complex (MHC) class II molecules, there are also environmental components suspected to contribute to the increased incidence of T1D. Among them are enterovirus, rotavirus, mumps virus or cytomegalovirus infections [3]; reduced levels of sphingolipids and sulfatides [4]; intestinal microbiota leakiness resulting in lower levels of butyrate [5]; changes in intestinal permeability in association with certain diets [6]; and beta cell stressors leading to the generation of neoepitopes [7] – all of which promote inflammation and increase the autoimmune propensity. While determining the specific etiology(s) of T1D will undoubtedly lead to new therapeutics aimed at preventing it entirely or at least halting its progression, biomarkers can also help in its early detection to guide the more-timely implementation of such therapeutics and significantly improve their efficacy. Therefore, there is a major interest in identifying reliable, predictive early biomarkers of T1D, which strikingly remain missing to date despite considerable efforts in this space.

Biomarkers can be direct cell products (e.g., DNA/RNA, proteins, and antibodies) and byproducts of enzymatic and metabolic processes (e.g., metabolites, peptides, and lipids), or inherent genetic determinants associated with a higher disease-risk (e.g., specific HLA haplotypes and gene polymorphisms). Current accepted biomarkers of T1D are based on genetic susceptibility in association with certain HLA haplotypes (e.g., DRB1 and DQB1) and the presence of antibodies against self-beta cell-antigens (e.g., GADA and IA-2A autoantibodies), as well as age, body mass index (BMI), and metabolic data derived from oral glucose tolerance tests (OGTT) [8-10]. While informative, such biomarkers have low predictive value [11, 12]. For example, islet autoantibodies (AAb) develop in ∼90% of individuals clinically diagnosed with T1D, but only ∼10% of children with one autoantibody progress to symptomatic T1D [13]. While individuals with multiple AAb are considered at a higher risk of progression, the risk level varies depending on the type and combination of the AAb among other risk factors, and some individuals with multiple AAb do not progress to clinical T1D diagnosis [14]. Thus, while highly associated with T1D diagnosis in retrospect, autoantibodies have limited utility in distinguishing non-progressors from progressors (i.e., those who develop symptoms and ultimately receive clinical diagnosis). Other biomarkers based on glucose metabolism like glycated hemoglobin A1C (HbA1C), fructosamine, glycated albumin, or 1,5-anhydroglucitol, also manifest too late in the T1D pathogenic process, further diminishing their value in early diagnosis. Therefore, complementing the existing biomarker repertoire with islet-and immune cell-specific biomarkers, and combining them into composite signatures across multi-omics domains could help in establishing useful predictors for various T1D risk levels and disease stages, including the early stages. Ultimately, this could improve our ability to identify progressors from non-progressors. Notably, such biomarkers will also contribute to our understanding of the disease pathogenesis and will shed light onto its heterogeneity. This knowledge will in turn introduce new possibilities for more efficient interventions for the prevention and treatment of T1D.

Despite challenges in identifying early biomarkers of T1D so far, the hope has not abated as there are success stories in other conditions where metabolism-based biomarkers have been identified to predict with acceptable levels of certainty the imminent disease manifestation or progression (e.g., diabetic retinopathy and kidney disease) [15-18]. Notably for T1D, cellular metabolism is crucial for the survival and function of the pancreatic islets and immune cells. Immunometabolism and metabolic programming of immune cells is responsible for their pro-inflammatory polarization and the autoimmunity against the pancreatic beta cells [19, 20]. Metabolic stress of the islet beta cells themselves also contributes to their own demise during T1D development [21-25]. Therefore, a better understanding of the changes in the metabolomes of islets and immune cells as well as their proteomes, lipidomes, and transcriptomes is likely to contribute a more comprehensive understanding of these disturbances during T1D development. This significant novel insight into the associated pathogenic mechanisms will also help generate reliable composite biomarker signatures of the early disease stages before significant and irreversible damage to the insulin producing beta cell mass has occurred. Various single-omics studies have identified biomarkers of beta cell stress and T1D pathogenesis, such as the circulating GAD65 protein, proinsulin to C-peptide ratio, miR-375 or the beta cell extracellular vesicles (EV) miR-21-5p, and differentially methylated DNA species of certain genes like *INS* or *IAPP* [26-30]. However, the predictive value of single biomarkers has proven limited in identifying progressors from non-progressors, likely due to the fragmented nature of information obtained that way.

We recently introduced the concept of integrated parallel multi-omics to provide a more inclusive view of disturbances occurring simultaneously in various systems of the same individuals who are considered at a high risk for developing T1D and to ultimately extract reliable composite biomarker signatures of the disease [31]. Despite the limited sample cohort in our studies, the integrated analyses of parallel metabolomics, proteomics, lipidomics, and transcriptomics data indicated increased activation and proliferation of specific immune cell subsets in T1D high-risk subjects compared to healthy controls. They also highlighted affected signaling pathways and provided associated biomarker candidates with demonstrated links to T1D. More recently, to overcome the limitations in the size of our relatively small cohort of samples, we explored a novel computational approach of data augmentation of the quadra-omics dataset to further enhance the ability to identify candidate T1D biomarkers for downstream validation [32]. We now expand on this previous work to validate selected candidates based on the comparison of their proteomics profile in the expansive TEDDY (The Environmental Determinants of Diabetes in the Young; https://teddy.epi.usf.edu/research/) and DAISY (Diabetes Autoimmunity Study in the Young; https://medschool.cuanschutz.edu/barbara-davis-center-for-diabetes/research/clinical-epidemiology/daisy-research) cohorts and on results from direct ELISA measurements performed in the original plasma samples utilized in the parallel quadra-omics studies [31, 32].

## RESULTS

### Selection of Biomarker Candidates Predicted in Augmented Parallel Dual-Omics Data for Further Validation

Our recent studies showed enhanced power of the Ingenuity Pathway Analysis (*IPA*)’s biomarker prediction algorithms in the augmented parallel multi-omics data compared to the original dataset from our limited cohort of nine samples from healthy, high-risk, and new-onset subjects. The quadra-omics data from these samples were imputed and amplified 1000 times to the equivalent of 9000 virtual subjects using proprietary algorithms, as previously described in detail [32]. In total, 41 candidate biomarkers were predicted by *IPA* in the augmented proteomics and metabolomics datasets compared to 19 in the corresponding original data obtained in samples from three subjects who were AAb positive and considered at a high risk of developing T1D. Of these candidates, 31 and 15 respectively had confirmed links to T1D based on our own review of published studies (Table 1).

**Table 1.** Candidate biomarkers predicted by *IPA* in the original and augmented proteomics and metabolomics datasets of subjects at high risk for developing T1D. Candidate biomarkers with a single underline were exclusively predicted in the augmented dataset. Candidates with double underline were predicted in common between the augmented and original datasets. All biomarker candidates have been implicated in T1D based on the published studies cited next to each.

| Original | Augmented |
| --- | --- |
| Biomarker name<sup>1</sup> | Biomarker name<sup>1</sup> |
| APOA2 [33], APOE [34], CD44 [35], <u>CETP</u> [36], GPNMB [37], <u>IGF1</u> [38], IGFBP2 [39], JAG1 [40], LDLR [41], MEP1B [42], MMP2 [43], <u>MMP9</u> [43], <u>PTPRC</u> [44], SELL [45], <u>VCAM1</u> [46] | <u>ACE</u> [47], <u>APOA1</u> [33], <u>APOB</u> [48], <u>APOC1</u> [49], <u>CASP3</u> [50], <u>CCL5</u> [51], <u>CD36</u> [52, 53], <u>CETP</u> [36], <u>CR2</u> [54], <u>CXCL12</u> [55], <u>FADD</u> [56, 57], <u>FAS</u> [56, 58], <u>FGFR1</u> [59], <u>GFAP</u> [60], <u>HPSE</u> [61], <u>HSPB1</u> [62], <u>IGF1</u> [38], <u>IGF2</u> [38], <u>IGHM</u> [63], <u>IL18</u> [64], <u>MASP2</u> [65], <u>MMP9</u> [43], <u>MSTN</u> [66], <u>PCSK9</u> [67], <u>PDE5A</u> [68], <u>PON1</u> [69], <u>PTPRC</u> [44], <u>RETN</u> [70], <u>SOD1</u> [71], <u>SRC</u> [72], <u>VCAM1</u> [46] |

<sup>1</sup> Biomarker names correspond to the associated gene names. Biomarker prediction was performed in the proteomics and metabolomics datasets independently and candidates were then consolidated since *IPA* cannot currently perform biomarker predictions in integrated multi-omics data.

Of the 31 candidate biomarkers predicted in the augmented data, 26 were unique and only 5 were also identified in the original dataset. Several of the unique candidates belonged to similar families or sub-families of proteins involved in metabolism, oxidative stress, immune function, and diabetes. We further examined the potential functional interconnectivity of these 26 candidates and their involvement in diseases and related biological functions using *IPA*’s knowledge base, which is based on scientific literature indexed in PubMed and other well recognized data repositories [73]. All candidates had direct connections with diabetes in general with a high significance (*p* < 0.000001 by Fisher’s test) and many were simultaneously implicated in lipid/lipoprotein metabolism, immune function, and inflammation (Figure 1).

**Figure 1.**
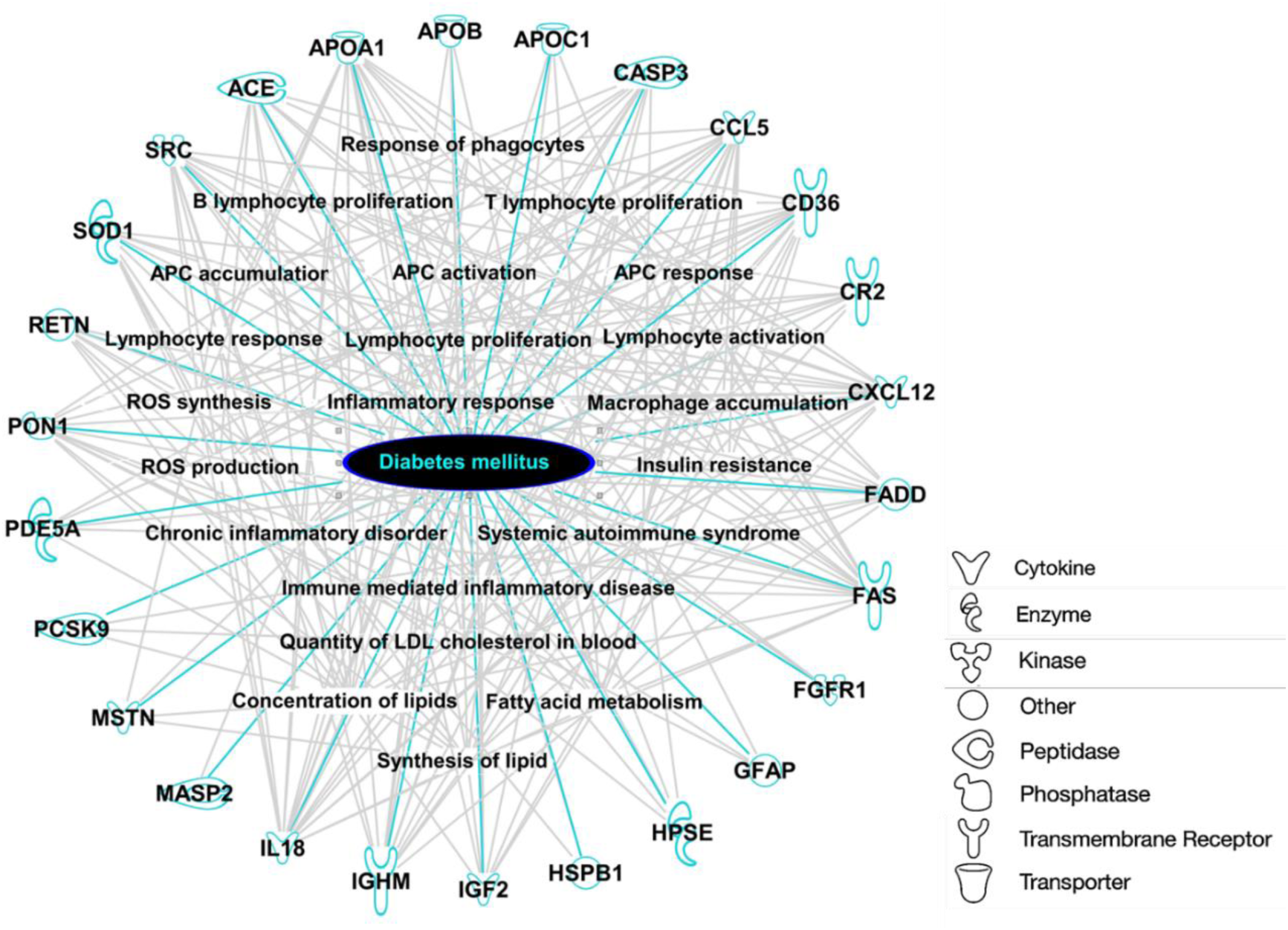
Overlay representation of genes (proteins) implicated in diabetes and various metabolic and immune/inflammatory functions, which were uniquely predicted by *IPA* as candidate biomarkers in the augmented dual-omics (proteomics and metabolomics) datasets obtained from the plasma of subjects considered at high risk for developing T1D. This representation highlights the interconnectivity of the candidate biomarkers and their impact on the various, yet related inflammatory diseases and biological functions simultaneously. All candidates had established connections to diabetes as highlighted in cyan color and their connections to additional inflammatory diseases and biological functions are shown in grey. The symbols associated with each gene represent the function of the corresponding protein as shown in the symbol key.

The 26 biomarker candidates uniquely predicted in the augmented dataset were narrowed down to 20 based on their confirmed detection in the untargeted proteomics analyses in blood samples of the TEDDY and DAISY cohorts [71, 74]. Then, further selection from the various represented protein families was made based on the change in their expression level in the high-risk subjects relative to the healthy controls. A threshold of ≥ 20% up or down change from the original proteomics data was used in this selection. This yielded 10 biomarker candidates, specifically: apolipoprotein C1 (ApoC1), angiotensin-converting enzyme (ACE), cluster of differentiation 36 (CD36; also known as scavenger receptor class B member 3 [SCARB3]), complement receptor 2 (CR2; CD21), fibroblast growth factor receptor 1 (FGFR1; CD331), immunoglobulin heavy constant mu (IGHM), proprotein convertase subtilisin/kexin type 9 (PCSK9), serum paraoxonase and arylesterase 1 (PON1), superoxide dismutase 1 (SOD1; SODC), and vascular cell adhesion protein 1 (VCAM1, CD106). Two additional candidates predicted in common between the augmented and original parallel multi-omics data and with strong established involvement in lipoprotein metabolism and T1D were also selected by the same criteria: cholesteryl ester transfer protein (CETP) and protein tyrosine phosphatase receptor type C (PTPRC; also known as CD45, CD45R, or B220). These 12 selected biomarker candidates were then all measured by ELISA in the samples originally utilized in the prior parallel quadra-omics studies for further validation (see Figure 4).

**Figure 2.**
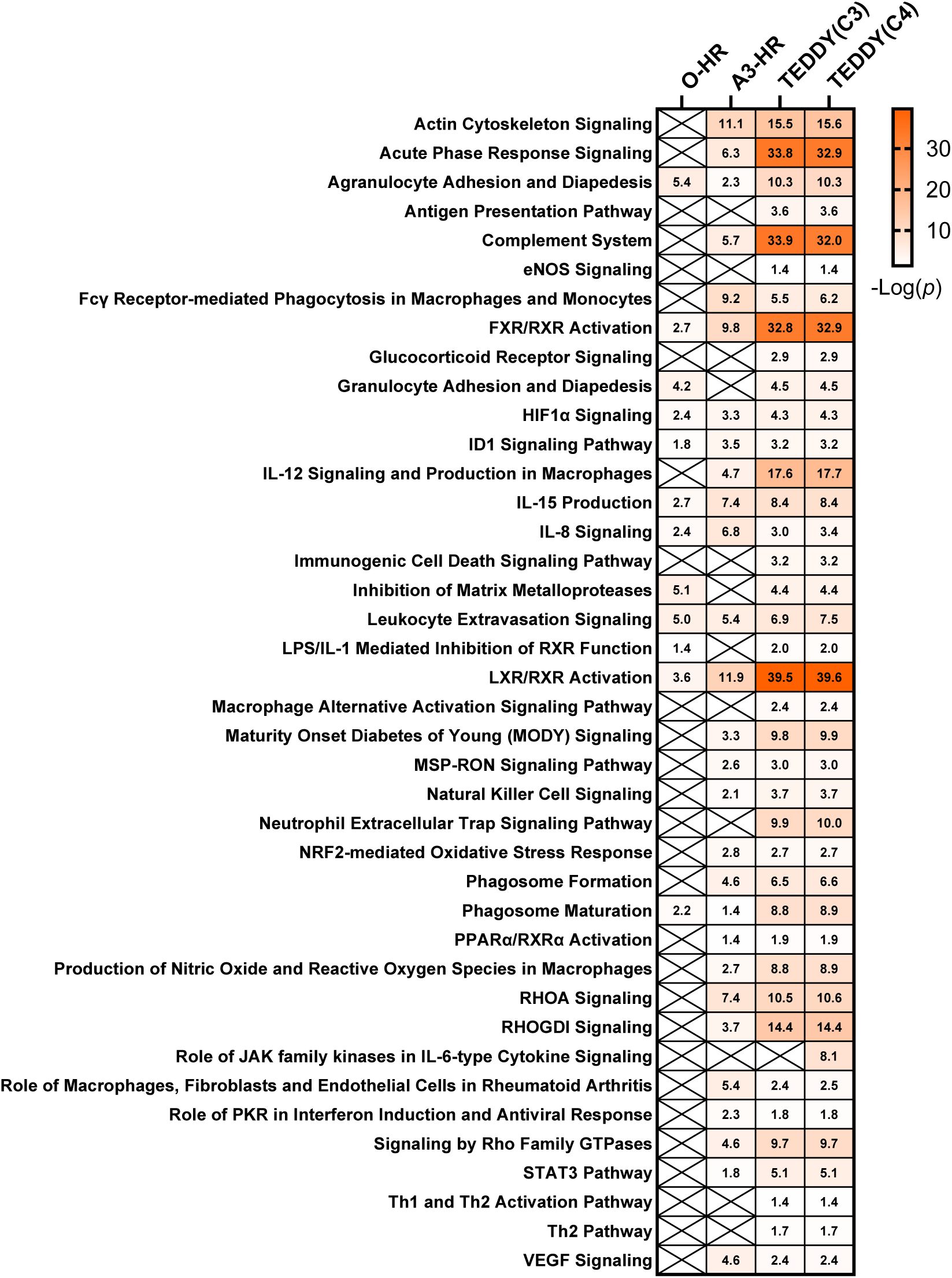
Heatmap showing canonical pathways identified by *IPA* in proteomics datasets obtained in our small cohort of high-risk subjects in the original (O-HR) and augmented (A3-HR) data as well as in the proteomics datasets obtained in the TEDDY cohort from seroconverted subjects that either progressed (C3; progressors) or did not progress to T1D diagnosis (C4; non-progressors); the C3 and C4 groups were also named T2 and I2, respectively, in reference [74]. The escalating color gradient in the boxes indicates increased significance of prediction in each dataset with corresponding −Log(*p*) values shown. Values ≥ 1.3 indicate significance (i.e., *p* ≤ 0.05 by Fisher’s test). Predictions with −Log(*p*) < 1.3 were not shown. Boxes marked with “X” indicate no prediction.

**Figure 3.**
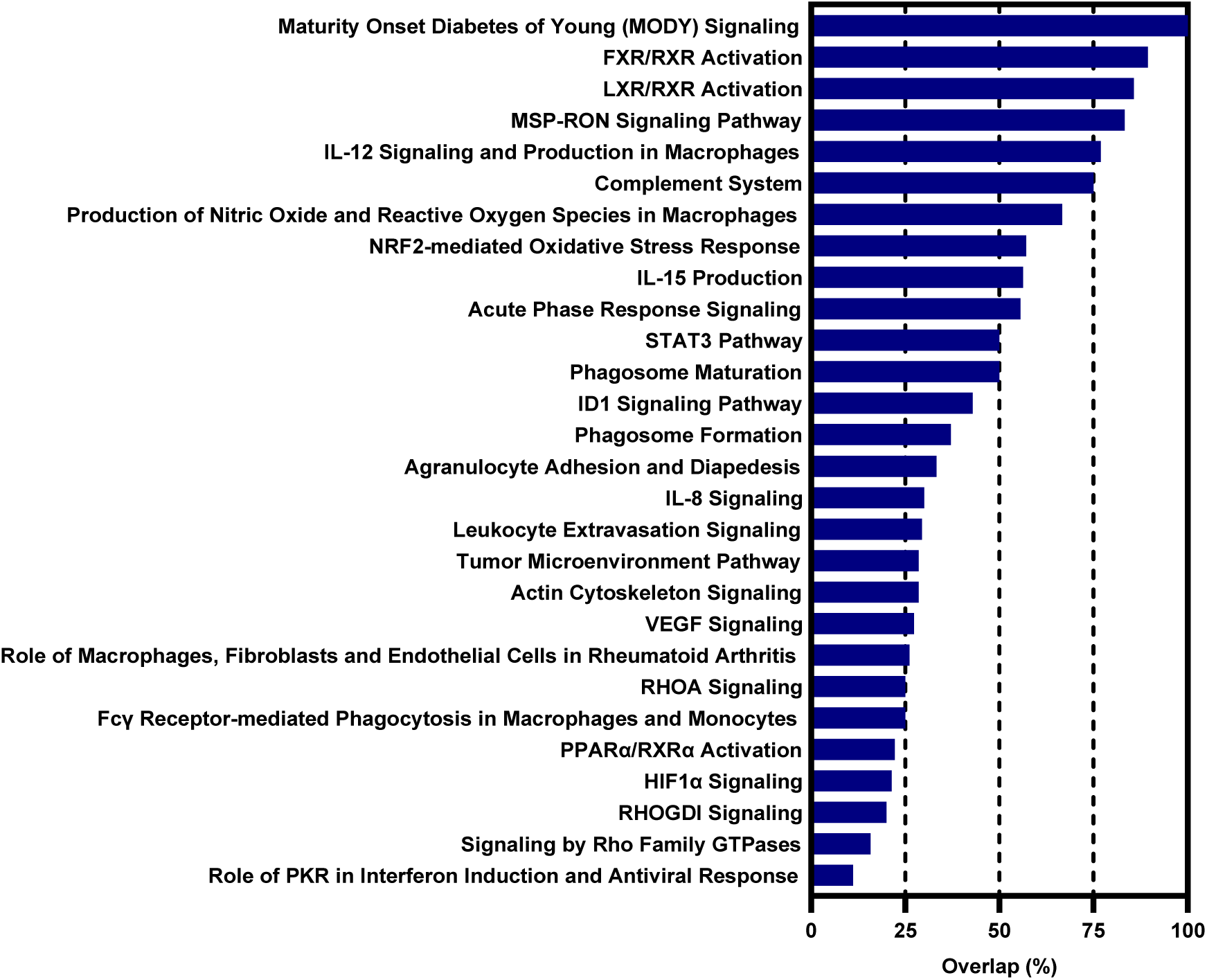
Bar graph showing the percent (%) overlap in features (proteins) associated with the shown immune/inflammatory canonical pathways that were identified in common between our augmented proteomics dataset from the high-risk subjects (A3-HR) [32] and the untargeted proteomics data obtained from seroconverted/non-progressor subjects (C4 = I2) of the TEDDY cohort [74].

**Figure 4.**
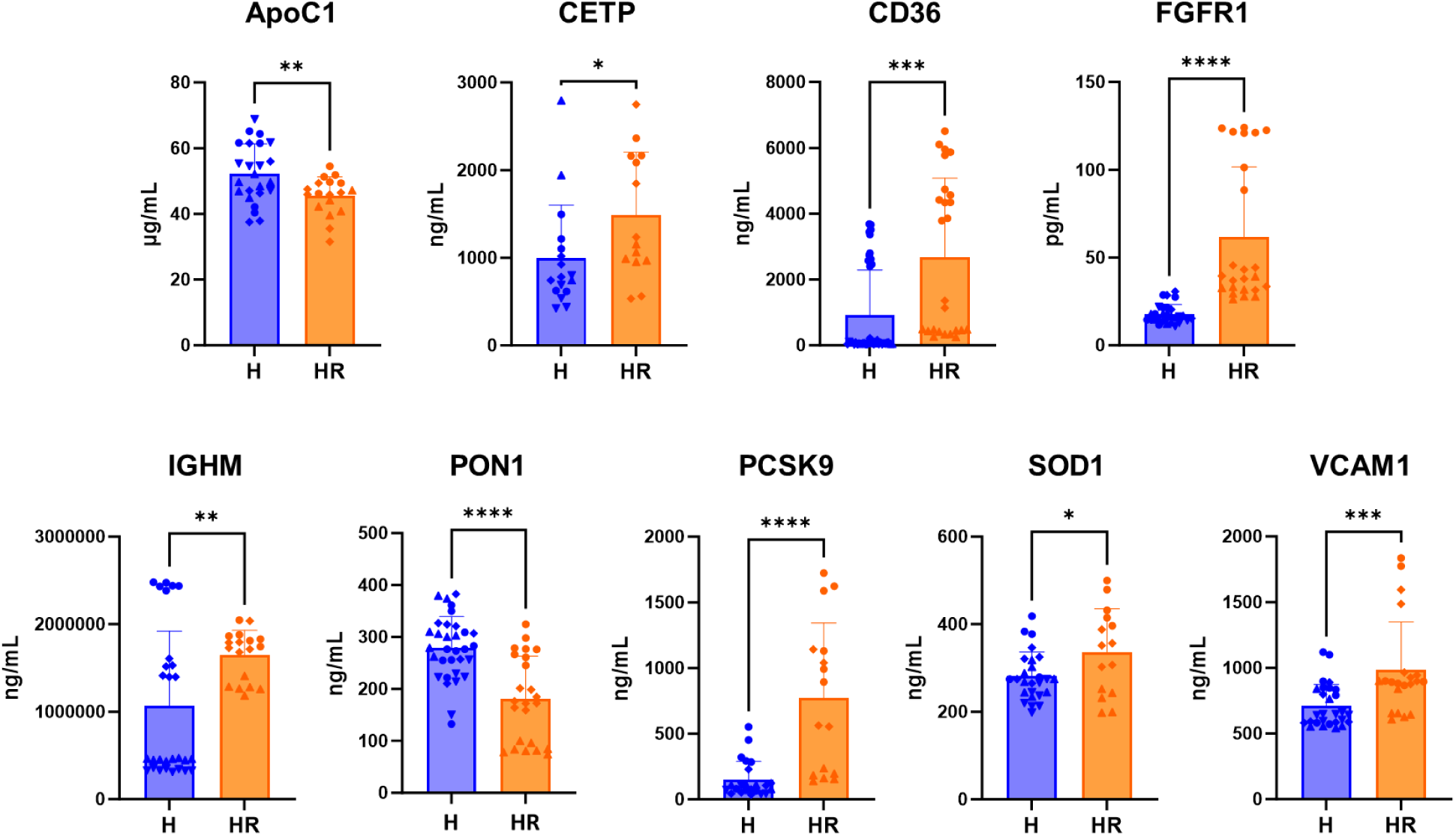
Protein levels of 9 of the 12 selected candidate biomarkers measured by ELISA in the blood samples of high-risk (HR; *n* = 3) subjects compared to those of healthy controls (H; *n* = 4). The remaining 3 candidates that were unchanged are shown in supplementary Figure S1. To ensure reproducibility of the results, measurements were conducted in replicates at various dilutions and repeated on 2 – 5 separate occasions in aliquots of the individual samples from each subject group. Replicates from each subject were shown in unique symbols (i.e., circles, diamonds, and triangles). Data were shown as means ± standard deviation (SD). Pairwise statistical comparisons were by two-tailed unpaired *t*-test; *, **, ***, and **** indicate *p*-values ≤ 0.05, ≤ 0.01, ≤ 0.001, and ≤ 0.0001, respectively.

### Pathway Analysis in the Augmented Proteomics Data versus TEDDY

To validate the biological relevance of the biomarker candidates predicted by *IPA* in the augmented data obtained from the AAb positive high-risk subjects in our small cohort, we performed parallel canonical pathway analyses using independently the original (O-HR) and the augmented proteomics (A3-HR) datasets of the same high-risk subjects as well as the previously obtained proteomics data from the AAb positive progressor (C3) and non-progressor (C4) groups from the TEDDY cohort; also named T2 and I2 respectively in reference [74]. The analyses were performed using *IPA*’s *Canonical Pathway* module with a focus on immune inflammatory functions, as previously described in detail [31]. Each identified pathway in the various datasets was assigned by *IPA* a prediction significance value (−Log[*p*]) and the results were compiled and presented side-by-side in a heatmap showing the predicted pathways and their corresponding −Log[*p*] values to highlight agreement, or lack thereof, in the pathway prediction across the various datasets (Figure 2). Forty pathways were identified in the data from the TEDDY-C4 group, 39 from the TEDDY-C3 group, 28 in the augmented dataset (A3-HR), and 12 in the original data (O-HR) from our small cohort. The increased number of pathways matching those from the TEDDY groups identified in the augmented dataset (A3-HR), which was derived from the original three high-risk subjects, highlighted the improved analytical power of *IPA* to a level comparable to that in datasets obtained from the substantially larger TEDDY-C3 and C4 groups containing 94 and 401 subjects, respectively. A sizable number of these pathways was not predicted in the original data and the few identified ones had lower prediction significance compared to those identified in the augmented dataset (Figure 2).

Further examination of the specific proteins associated with the canonical pathways identified in common between the augmented data from the three high-risk subjects and the data from the seroconverted (AAb positive) non-progressor TEDDY-C4 (I2) group confirmed a high degree of overlap in such proteins (Figure 3). Out of the 28 identified pathways, 6 had ≥ 75% of overlap, 6 ≥ 50%, 11 ≥ 25%, and only 5 had < 25% of overlap in the proteins contributing to the prediction of the listed pathways (see supplementary Table S1 for comprehensive list of these proteins across all the subject groups). Together, these results indicated a significant unmasking by our data augmentation approach of biologically relevant proteins that were otherwise undetected in the original data from the limited number of samples.

### Validation of Candidate Biomarkers on the Protein Level

To further validate the 12 biomarker candidates identified through the data augmentation approach [32] and the selection process described above, we quantified their levels using ELISA-based methods in the original plasma samples of the high-risk and healthy subjects, which were previously utilized in the parallel quadra-omics studies [31]. The ELISA measurements showed markedly reduced (*p* < 0.05 by ANOVA) protein levels of ApoC1 and PON1, whereas CETP, CD36, FGFR1, IGHM, PCSK9, SOD1, and VCAM1 were elevated in the high-risk subjects compared to the healthy controls (Figure 4). ACE, CD45, and CR2 were unchanged (supplementary Figure S1).

### Glycated ApoC1 as a potential early biomarker of T1D

ApoC1 (apolipoprotein C1) is the primary inhibitor of CETP (cholesteryl ester transfer protein). The above ELISA measurements showed marked reduction in ApoC1 and a parallel increase in CETP levels in the plasma of the high-risk subjects compared to the healthy controls (Figure 4). Since CETP plays an essential role in determining the composition of high-density lipoproteins (HDL) and low-density lipoproteins (LDL) among other cholesterol species, we examined whether post-translational modification of ApoC1 through glycation further contributes to its functional impairment and reduces its inhibition of CETP, whereby impacting on the HDL/LDL cholesterol composition, as previously suggested [49]. We performed Western blot analysis to examine whether ApoC1 exhibited a smearing pattern consistent with a mass-shift associated with post-translational modification in the high-risk subjects but not in the healthy controls. However, the results showed a similar smearing pattern in both subject groups (Figure 5A). This mass-shift was not observed for the purified recombinant protein. Consistent with the ELISA measurements, estimation of the ApoC1 levels from this semi-quantitative analysis suggested lower ApoC1 plasma levels in the high-risk subjects compared to the healthy controls (Figure 5B). However, we found no differences in the various cholesterol species (e.g., HDL, LDL, and VLDL) measured in the same plasma samples from both subject groups of this small cohort (Figure 5C), the same plasma samples utilized in the Western blot analysis and the ELISA measurements.

**Figure 5.**
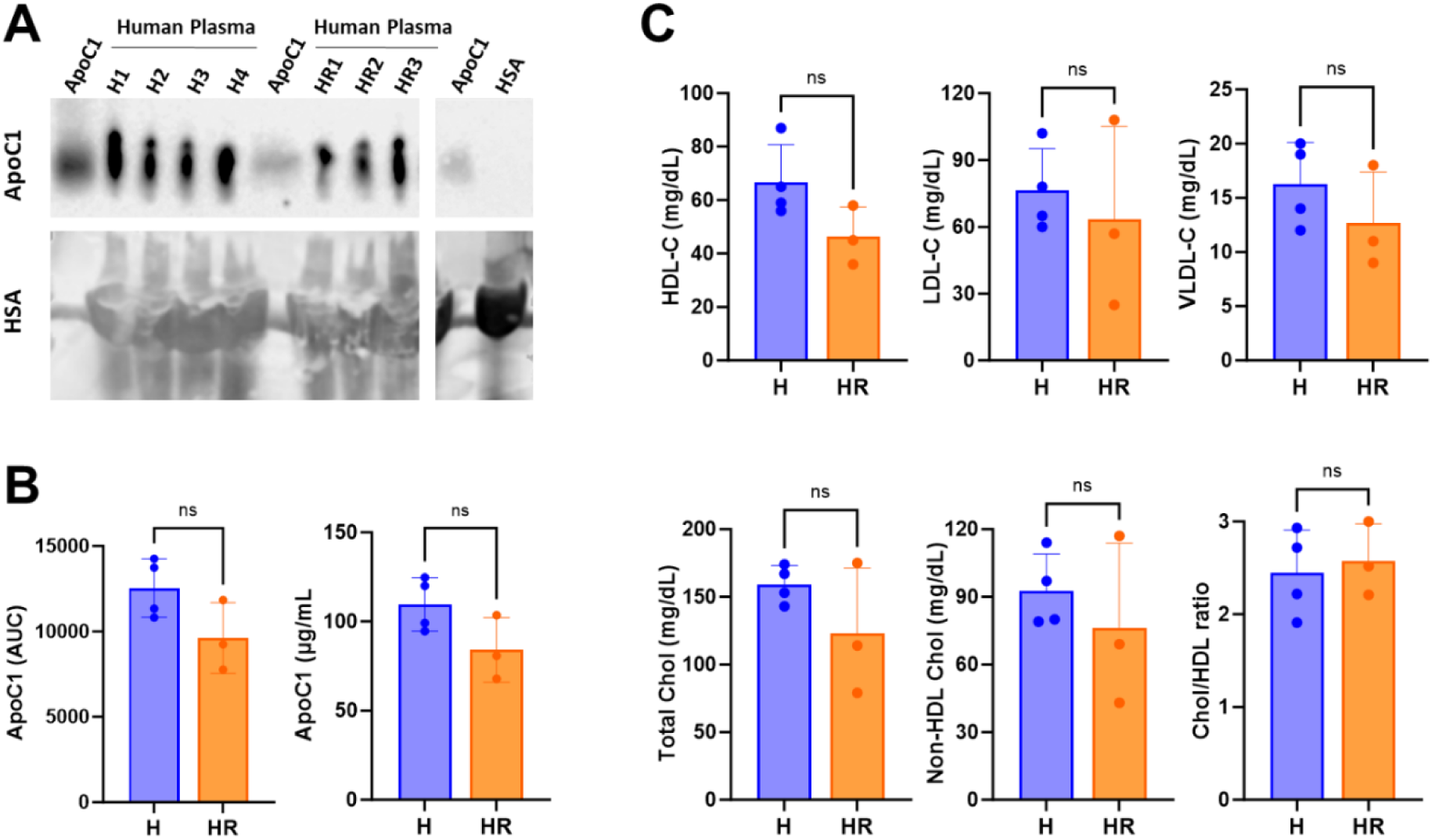
(A,B) Western blot analysis of ApoC1 in plasma of the healthy (H; *n* = 4) and high-risk (HR; *n* = 3) subjects. (A) shown on the top is ApoC1 and on the bottom human serum albumin (HSA). Purified recombinant ApoC1 protein and HSA were used as positive controls. The far-right control lanes are part of the same gel excluding irrelevant intermediary lanes. Given the high abundance of HSA in the samples, its bands were large and overlapped in the adjacent lanes due to overloading, which could not be avoided by further dilution of the samples as that lead to losing the ApoC1 bands; depleting it from the samples was not attempted either for the same reason. (B) Semi-quantitative analysis of ApoC1 plasma levels shown as area under the curve (AUC) and as concentration (µg/ml) estimates based on the control recombinant ApoC1. (C) Measurements of various cholesterol (Chol) species in the plasma of the same subject groups (H, *n* = 4; HR, *n* = 3). Measurements were acquired using a Roche Cobas 6000 Chemistry Analyzer (Roche Diagnostics; Indianapolis, IL, USA). Data in B and C were shown as means ± standard deviation (SD). Pairwise statistical comparisons were by unpaired *t*-test; ‘ns’ indicates *p*-value > 0.05.

## DISCUSSION

The goal of the current work is to continue building on our progress in the T1D biomarker space by validating candidate biomarkers previously identified in our studies employing *(a)* parallel multi-omics to obtain more inclusive data on metabolic, genetic, and immunologic disturbances in individuals at high risk for developing T1D [31] and *(b)* a computational data augmentation approach to address the inherent limitations of small input data obtainable from precious biological samples that are, not only difficult to collect from vulnerable populations like children, but also very costly to obtain on a large scale [32]. These logistical issues remain a significant challenge in this type of studies. Nonetheless, despite the small cohort of samples, our prior studies resulted in a promising list of candidate biomarkers, which we aimed to validate here and integrate into a potential composite signature of T1D risk in those subjects who are considered at a high risk for developing the disease according to the TrialNet classification [75]. Our long-term goal is to perform this type of studies in larger cohorts of longitudinal samples from at-risk subjects during their disease progression to ultimately identify composite biomarker signatures of the various disease stages and be able to potentially discriminate progressors from non-progressors. Future studies will also examine the correlation of these disease-stage specific biomarker signatures with clinical data on metabolic changes (e.g., fasting glycemia and HbA1C) and peripheral immune phenotypes and cytokine profiles during the progression of the disease.

As a first step in the validation process, we selected 12 from the 41 candidate biomarkers that were predicted in the augmented proteomics and metabolomics datasets from the high-risk subjects (Table 1). Selection was by cross-reference to a similar group of 401 subjects in the TEDDY cohort who were seroconverted (AAb positive) but without clinical disease at the time of analysis (i.e., non-progressors) [76] and by other ranking criteria as detailed above. We next performed protein measurements by ELISA in the original plasma samples where the quadra-omics studies and the biomarker prediction were performed based on LC-MS/MS assessments (Figure 4) [31]. To ensure reproducibility of the results in the current limited number of samples in each subject group of the original cohort, the ELISA measurements were performed in replicates with multiple repeats at different times using serial sample dilutions. These measurements showed marked differences (*p* ≤ 0.05) between the T1D high-risk subjects and the healthy controls in all but 3 of the 12 selected biomarkers, namely, ACE, CD45, and CR2 (supplementary Figure S1). The remaining 9 candidates either increased or decreased in the high-risk subjects relative to the healthy controls. Specifically, CETP, CD36, FGFR1, IGHM, PCSK9, SOD1, and VCAM1 were elevated, whereas ApoC1 and PON1 were reduced (Figure 4). Noticeably, all these candidate biomarkers had established connections with lipid/lipoprotein metabolism, immune function, inflammation, and diabetes (Figure 1). Their functions, potential contributions to inflammation, autoimmunity, and involvement in T1D pathology as well as their possible utility in a potential composite signature of T1D risk are discussed briefly below.

ApoC1 is the primary inhibitor of CETP, which plays an essential role in plasma lipoprotein and cholesterol (HDL/LDL) composition by mediating the transfer and exchange of cholesteryl ester and triglycerides on plasma proteins. It was previously suggested that post-translational modification of ApoC1 by glycation impairs its inhibition of CETP, which may be the reason for the abnormal plasma lipoprotein composition in diabetic patients [49, 77]. Since the ApoC1 levels were reduced and those of CETP were elevated (*p* < 0.05) in the high-risk subjects, we tested the hypothesis that post-translational modification of ApoC1 (possibly through glycation or glycosylation) could serve as an early T1D biomarker in the high-risk subjects. The Western blot analysis, however, showed similar smearing patterns in the ApoC1 band across the samples, suggesting similar post-translational modification in all subject groups, modification which was not observed in the purified recombinant protein (Figure 5A). We attempted to check whether glycosylation contributed, at least in part, to the post-translational modification of ApoC1 by treating all the plasma samples with glycosidases prior to the Western blotting; however, these attempts failed due to the complete loss of the ApoC1 band in the treated samples (supplementary Figure S2). Further examination of glycated peptides in our LC-MS/MS proteomics data revealed apolipoproteins ApoB and ApoH to be glycated, but not ApoC1 (supplementary Table S2). Therefore, we were unable to establish the value of glycated ApoC1 as an early T1D biomarker in addition to its reduced total protein levels in the high-risk subjects. Similarly, PON1 was also reduced (*p* < 0.05) in the high-risk subjects as determined by the ELISA measurements. PON1 is a plasma paraoxonase originally discovered for its hydrolytic and detoxifying activity of organophosphate pesticides. PON1 is a powerful antioxidant of lipid peroxides on LDL cholesterol, and its homocysteine thiolactonase activity prevents the *N*-homocysteinylation of proteins and preserves their structure and function [78]. Since ApoC1, CETP, and PON1 are intimately involved with plasma lipoproteins, we further examined the composition of HDL and LDL among other cholesterol species in the plasma of the subject groups in our small cohort. We did not find significant differences in the cholesterol profiles, likely due to the limited cohort of samples; although, a mild trend towards lower HDL was noticeable in the high-risk subjects (Figure 5C). Future studies in larger cohorts are needed to examine this potential correlation of lower ApoC1 and PON1 with higher CETP and an associated skewing in the HDL/LDL composition in at-risk subjects in comparison to healthy individuals, as has been previously shown for diabetic patients [77]. Nevertheless, the ELISA measurements showing, on the protein level, markedly reduced ApoC1 and PON1 and elevated CETP supported their potential utility in a composite signature of T1D risk. Notably, PON1 and ApoC1 were used as biomarkers of dyslipidemia (NCT00903617) and myocardial infarction [79], but neither had been previously suggested as a biomarker of autoimmune T1D. Thus, the current findings further highlight the promise of our computational approach of data augmentation in multi-omics datasets in the identification of novel biomarkers of T1D risk.

As noted above, CD36, FGFR1, IGHM, PCSK9, SOD1, and VCAM1 were elevated in the high-risk subjects compared to healthy controls (*p* < 0.05). Like ApoC1 and PON1, CD36 and PCSK9 are also involved in lipid metabolism and the processing of HDL/LDL cholesterol. CD36 is a membrane glycoprotein expressed on the surface of many tissues and cell types (e.g., endothelial cells, adipocytes, and macrophages). It binds several ligands and has been implicated in various physiological functions and pathological conditions (e.g., hemostasis and atherogenesis). As a member of the class B scavenger receptor family and a fatty acid translocase, CD36 binds free fatty acids and oxidized LDL cholesterol and facilitates their update into cells. In immune cells, binding of oxidized LDL triggers pro-inflammatory signaling cascades attributed to atherogenesis [80]. Elevated levels of circulating soluble CD36 have also been implicated in type 2 diabetes (T2D) [81]; but a recent study by Castelblanco and colleagues showed no significant differences in patients with either T1D or T2D compared to non-diabetic subjects [82]. Our ELISA measurements showed a significant increase (*p* = 0.0003) in those considered at a high risk of developing T1D compared to healthy controls (Figure 4). Also, PCSK9 was increased in the high-risk subjects (*p* < 0.05). PCSK9 is a member of the proprotein convertase family, and it too is expressed in many tissues and cell types. PCSK9 plays a major role in the homeostasis of LDL cholesterol by regulating its receptor (LDLR). A major pathway of LDL uptake by cells is through binding to LDLR. Upon internalization of the LDL-LDLR complex, LDLR is released in the endosome and recycled back to the plasma membrane for additional rounds, which results in the lowering of extracellular LDL cholesterol. PCSK9 also binds to LDLR; however, it prevents recycling of LDLR to the plasma membrane and directs it instead to the lysosome for degradation, whereby resulting in higher levels of extracellular and circulating LDL cholesterol. Because of this, PCSK9 inhibitors have gained significant attention in recent years as LDL-lowering drugs [83, 84]. Moreover, it has been shown that loss-of-function mutations in the *PCSK9* gene result in lower fasting glycemia and LDL, suggesting that elevated PCSK9 could have the opposite effect [85]; although, we did not find significant changes in the HDL/LDL composition in our cohort, which was likely due to the current small sample size. Nonetheless, the consistent pattern we observed in ApoC1, CETP, PON1, CD36, and PSCK9, all of which have intimate involvement in lipid cholesterol metabolism, which is known to be disrupted in T1D, support their potential utility in a composite biomarker signature of increased risk for developing T1D.

Moreover, the other candidate biomarkers identified here FGFR1, IGHM, SOD1, and VCAM1 have no direct links to lipid cholesterol metabolism, but they are involved in inflammation, autoimmunity, and diabetes (Figures 1 and 2), and they were all markedly elevated in the high-risk subjects (*p* < 0.05). FGFR1 is a membrane receptor with tyrosine kinase activity found on the surface of cells. It activates different downstream signaling pathways depending on the type of cell and the fibroblast growth factor ligand that binds to it (e.g., FGF1 – 10). Mutations in the *FGFR1* gene are implicated in various congenital abnormalities of the musculoskeletal system as well as the developing pancreas [86]. FGFR1 is also implicated in carcinogenesis, where its gain-of-function mutations have been reported in multiple types of cancer [87, 88]. Also, mice with dominant-negative forms of FGFR1 have reduced pancreatic beta cell mass and disturbed glucose homeostasis, and they spontaneously develop diabetes [89]. A previous MALDI–TOF proteomics study by Chen and colleagues [59] showed an approximate 1.5-fold decrease in plasma FGFR1 in T1D patients when compared to healthy subjects and our ELISA measurements consistently showed a marked increase in the high-risk subjects. Hence, the cumulative evidence points to involvement of FGFR1 in T1D pathogenesis. Moreover, the ELISA measurements also showed a similar pattern of IGHM in the same subjects. IGHM is a part of the B cell receptor and plays a crucial role in the early development of B lymphocytes and their subsequent negative selection against self-antigens and further functional maturation towards antibody producing B lymphocytes [90, 91]. IGHM has been implicated in T1D development in the non-obese diabetic (NOD) mice through preferential antigen presentation of pancreatic beta-cell antigens by the B lymphocytes, whereby contributing to the loss of self-tolerance and progression of anti-beta cell autoimmunity [92, 93]. Thus, the higher levels of IGHM in the AAb positive high-risk subjects are consistent with the increased potential of anti-beta cell autoimmunity and T1D development. Notably, Chen and colleagues found a 1.34-fold increase in plasma IGHM of T1D patients relative to the healthy subjects in their MALDI–TOF proteomics study [59], whereas our LC-MS/MS proteomics found no differential expression of IGHM among the three subject groups in our relatively small sample cohort [31]. This difference likely resulted from the variation between the two proteomics approaches and the ELISA measurements. Indeed, while ELISA measurements here confirmed the trend in the protein levels measured earlier by LC-MS/MS proteomics in 6 of the 9 selected candidate biomarkers (positive slopes), there was only limited correlation (supplementary Figure S3), which is again likely due to the small cohort and inherent differences of the analytical methods. Nevertheless, the elevated protein levels of FGFR1 and IGHM as measured by ELISA in the high-risk subjects support their utility as biomarkers in a potential composite signature of T1D risk. Remarkably, despite the limited correlation between the methodologies, our computational approach of omics data augmentation resulted in meaningful candidate biomarkers that were not identified in the small cohort–derived original data.

SOD1 (superoxide dismutase 1) is a powerful anti-oxidant that was also elevated (*p* < 0.05) in our ELISA measurements in plasma of the high-risk T1D subjects (Figure 4). This finding is consistent with previous longitudinal proteomics and ELISA analyses conducted by Liu and colleagues during the natural progression of T1D in at-risk individuals, before and after seroconversion and leading up to their clinical diagnosis [71]. Therefore, SOD1 may be an important biomarker in a potential composite signature of T1D risk.

Lastly, the pro-inflammatory vascular cell adhesion protein 1 (VCAM1) was also elevated (*p* < 0.05) in the plasma of the high-risk subjects compared to both the new-onset patient and healthy controls (Figure 4). VCAM1 is a member of the immunoglobulin superfamily that is primarily expressed on vascular endothelial cells. Its expression is upregulated by pro-inflammatory T helper 1 (Th1) cytokines like IL-1b and TNF-α. VCAM1 binds to its cognate integrin α4β1 (also known as Very Late Antigen-4, VLA-4) on circulating leukocytes to facilitate their transendothelial migration and extravasation into the interstitial space and recruitment to inflamed tissues [94]. VCAM1 is also cleaved from the plasma membrane by the metalloprotease 17 (ADAM17; also known as tumor necrosis factor-α-converting enzyme, TACE), and the circulating soluble form is considered a biomarker of inflammation in various conditions like atherosclerosis, age-associated neurodegeneration, and cancer progression and metastasis [95, 96]. In T1D, it was shown that VCAM1 plasma levels (measured by ELISA) were unchanged in children with established T1D compared to age-matched healthy controls [97]. Our own ELISA measurements showed higher VCAM1 levels in the high-risk subjects. Therefore, VCAM1 may also prove of value as an early biomarker in a potential composite signature of T1D risk, among the other above biomarkers.

## CONCLUSSIONS

In summary, the findings in the present study further highlight the benefits of both our parallel multi-omics [31] and the more recently described data augmentation approach [32] in the unmasking of important patterns and features, which are otherwise undetectable in small biological datasets derived from limited cohorts, to identify promising candidate biomarkers of T1D for downstream validation. The current analyses also demonstrate the biological relevance of ApoC1, CETP, CD36, FGFR1, IGHM, PON1, PCSK9, SOD1, and VCAM1 in the context of the expansive TEDDY cohort and support their utility as biomarkers in a potential composite signature of T1D risk. While we continue to investigate new methods of data integration and exploration of parallel multi-omics datasets from constantly expanded cohorts to grow the list of validated T1D biomarkers, we hope that the current signature of reduced ApoC1 and PON1 and elevated CETP, CD36, FGFR1, IGHM, PCSK9, SOD1, and VCAM1 will stand the test of time in clinical evaluations of its association with increased risk of developing T1D.

## MATERIALS AND METHODS

### Human Plasma Samples and Multi-Omics Analyses

Blood samples were collected as previously described [31] in purple top (EDTA) Vacutainers (BD Biosciences; Franklin Lakes, NJ, USA) from subjects considered at a high risk of developing T1D (*n* = 4) during routine visits as part of the ongoing TrialNet’s Pathway to Prevention screening Study (TN-01) (https://www.trialnet.org/our-research/risk-screening) and from healthy subjects (*n* = 4) as part of another study approved by the IRB of the University of Miami (study number 11995-115). The classification of the high-risk subjects was based on the TrialNet classification [75]. One of the four high-risk subjects exhibited signs of abnormal oral glucose tolerance test (OGTT) and was confirmed to have converted to a new-onset patient during a second OGTT and another sample collection two weeks later (*n* = 2 samples). This subject was excluded from the analysis presented here. We did not have access to samples from low-risk subjects, as per the TrialNet classification [75], in the TN-01 study at the time of sample collection and the samples from the healthy subjects obtained under UM study number 11995-115 were used for all the comparisons. Plasma was obtained immediately after the blood collection and stored at –80 °C until analysis. The reader is referred to supplementary Table S1 in reference [31] for complete details of the samples and clinical data available on the subjects at the time of sample collection. Quadra-omics analyses (proteomics, metabolomics, lipidomics, and transcriptomics) were performed on aliquots of the same samples from each subject as previously described in detail [31].

### Proteomics Analyses

Proteomics analyses were performed in the TEDDY/DAISY cohorts and in the small cohort of samples noted above, as previously described in detail in [74, 98] and [31], respectively. In brief, LC-MS/MS analysis in the small cohort was done on a Waters NanoAquity UPLC system with a custom packed C18 column (70 cm × 75 μm i.d., Phenomenex Jupiter, 3 μm particle size, 300 Å pore size) coupled to a Q-Exactive mass spectrometer (Thermo Fisher Scientific). Peptides, in diluted samples depleted of the 14 most abundant plasma proteins (albumin, IgG, α1-antitrypsin, IgA, IgM, transferrin, haptoglobin, α1-acid glycoprotein, α2-macroglobulin, apolipoprotein A-I, apolipoprotein A-II, complement C3, transthyretin and fibrinogen), derivatized, and fractionated into 24 fractions by high-pH reversed phase chromatography as before [99], were eluted with a gradient of water (solvent A) and acetonitrile (solvent B). Analytes were analyzed online by nanoelectrospray ionization. Full mass scans were collected from 300 to 1800 *m/z* at a resolution of 35,000 at 400 *m/z*. Tandem mass spectra were collected in data-dependent acquisition of the top 12 most intense parent ions using high-energy collision induced dissociation (HCD) fragmentation (2.0 *m/z* isolation width; 30% normalized collision energy; 17,500 resolution at 400 *m/z*), before being dynamically excluded for 30s. A quality control sample, which consists of a protein digest of the bacterium *Shewanella oneidensis*, was also run before and after each batch of samples for additional quality control following a routine procedure [100]. Data were extracted with Decon2LS_V2 and DTA Refinery [101, 102] for mass recalibration and peak list extraction. Peptide identification was done with MSGF+ [103] using the human SwissProt database (downloaded from Uniprot Knowledgebase on February 22, 2019) with analysis parameters as previously defined [31]. Spectral-peptide matches, peptides and protein were filtered with MSGF probabilities to maintain false-discovery rate below 1%. The intensities of TMT reporter ions were extracted with MASIC [104] for quantitative analysis.

### Analysis of Multi-Omics Data and Identification of Candidate Biomarkers in Augmented Datasets

Analysis of augmented multi-omics data (proteomics, metabolomics, lipidomics, and transcriptomics) was performed using the *Biomarker Prediction* module of the *Ingenuity Pathway Analysis (IPA)* software package (Qiagen Bioinformatics; Redwood City, CA, USA; https://www.qiagenbioinformatics.com/products/ingenuity-pathway-analysis; RRID:SCR_008653) [73]). The original data containing 2292 proteins, 328 miRNAs, 75 metabolites, and 41 lipids, which were identified without exception in all nine analyzed samples, were augmented to the equivalent of 9000 virtual subjects, as was recently described in detail [32]. Briefly, 3 sets of augmented multi-omics datasets were previously generated at different imputation levels around the “golden ratio” of the data imputation (38.2%:61.8%) [105]. Here, we used the augmented dataset “A3” (with 40% imputation level), which yielded the highest number of predicted biomarker candidates [32]. The biomarker prediction analysis, which was performed on 4000 healthy and 3000 high-risk virtual subjects in A3, resulted at the time in 41 candidate biomarkers of diabetes mellitus (inclusive of T1D and T2D), 28 of which with confirmed direct links to T1D based on *IPA*’s knowledge database at the time of analysis. Since then, 3 other candidates (namely, ApoC1, CD36, and SOD1) have also been confirmed in the context of T1D based on the published literature, as noted in Table 1. Therefore, a total of 31 biomarker candidates were considered in this study, of which, 12 were selected for further investigation by ELISA protein measurements in the samples of high-risk and healthy subject groups in our small cohort previously used in the multi-omics analyses. Selection of the 12 candidates was primarily based on their detection in the TEDDY and/or DAISY clinical cohorts in addition to other criteria as described above in the results.

### Protein measurements by enzyme-linked immunosorbent assay (ELISA)

Protein levels of the selected 12 biomarker candidates were measured by ELISA in aliquots of the samples from each subject in the healthy and high-risk groups. The following ELISA kits were used for ApoC1 (Catalog # EHAPOC1), CR2 (Catalog # EH81RB), IGHM (Catalog # BMS2098), PCSK9 (Catalog # EH384RB), PON1 (Catalog # EH376RB), SOD1 (Catalog # BMS222), VCAM-1 (Catalog # BMS232), and CD36 (Catalog # EH88RB) from Thermo Fisher Scientific (Waltham, MA, USA); CETP (Catalog # NBP2-75239) from Novus Biologicals (Centennial, CO, USA); ACE (Cat # OKCD05600), CD45 (Catalog # OKEH05735-96W), and FGFR1 (Catalog # OKEH00124) from Aviva Systems Biology (San Diego, CA, USA). All assays were performed according to the manufacturers’ recommendations. Samples dilutions ranged between non-diluted and 1/100,000. All samples were measured in a minimum of triplicates per assay in a total of 2 – 5 repeats on separate occasions. Between group differences were analyzed by Student’s *t*-test (unpaired, two-tailed) using GraphPad Prism (GraphPad, La Jolla, CA).

### Evaluating ApoC1 post-translational modifications by Western blotting

To avoid jellification, 3 µL of each of the human plasma samples from the small cohort were diluted in 17 µL of Laemmli sample buffer (Bio-Rad; Hercules, CA, USA) followed by denaturing at 70 °C for 10 min and the resulting denatured 20 µL samples were loaded onto 12% Bis-Tris PAGE gel (GenScript; Piscataway, NJ, USA). 0.5 µg of purified recombinant ApoC1 (#CSB-EP001930Hue1; Cusabio, Houston, TX, USA) and 50 µg of human serum albumin (HSA; Human Albumin USP Albutein 25%; Grifols, Los Angeles, CA, USA) were loaded in separate parallel wells/lanes as controls. After electrophoresis (80 V for 3 h) using Tris-MOPS-SDS buffer (GenScript; Piscataway, NJ, USA), proteins were transferred onto a 0.2 µm pore size nitrocellulose membrane (Bio-Rad; Hercules, CA, USA) in a buffer (25 mM Tris, 192 mM glycine, 20% methanol, pH 8.3) using a constant current of either 100 mA for 20 minutes and 120 mA during 60 minutes, respectively, for the ApoC1 assessment and to ensure transfer based on HSA control. Membranes were then blocked in 2% fat-free milk in TBS solution containing Tween-20 (TBS-T) for 1 hour at room temperature. Blots were incubated overnight at 4 °C with mouse monoclonal anti-ApoC1 antibody (#sc-101263; Santa Cruz Biotech, Dallas, TX, USA) or mouse monoclonal anti-HSA antibody (#66051-1-Ig; ProteinTech; Rosemont, IL, USA). After incubation, membranes were washed three times with TBS-T, incubated with horse-radish peroxidase-linked recombinant mouse IgG Fc binding protein (#525409; Santa Cruz Biotech, Dallas, TX, USA) for 1 hour at room temperature, and finally washed three times in TBS-T. Detection of the immunoreactive bands was performed using the chemiluminescent reagent SuperSignal West Femto Maximum Sensitivity Substrate (Thermo Fisher Scientific; Waltham, MA, USA). Chemiluminescence images were acquired by Azure 600 Imaging System (Azure Biosystems; Dublin, CA, USA). ApoC1 bands were quantified by densitometry using *ImageJ* software version 1.54f (National Institutes of Health, Bethesda, MD, USA).

## SUPPLEMENTARY MATERIALS

Table S1: List of proteins contributing to the various degrees of overlap in the canonical pathways identified in the original (O-HR) and augmented (A3-HR) proteomics data from the small cohort and those from the C3 and C4 subject groups of the TEDDY cohort; Table S2: List of all peptides identified with post-translational modifications including glycation; Figure S1: ELISA measurements of selected candidate biomarkers that were unchanged between high-risk and healthy controls; Figure S2: Western blot analysis of ApoC1 glycosylation; Figure S3: Correlation analysis of ELISA and LC-MS/MS proteomics measurements.

## AUTHOR CONTRIBUTIONS

Conceptualization, M.H.A., P.B., and G.R.; Data curation, O.A., E.S.N., P.B., and M.H.A.; Formal analysis, O.A., S-T.C., G.R., M.O., B.W-R., E.S.N., P.B., and M.H.A.; Investigation, O.A., S-T.C., G.R. and E.S.N.; Software, G.R. and M.O; Funding acquisition, M.O, E.S.N., P.B. and M.H.A; Project administration, M.H.A. and P.B.; Supervision, P.B. and M.H.A.; Writing— original draft, O.A., P.B., and M.H.A.; Writing—review and editing, O.A., S-T.C., G.R., M.O., B.W-R., E.S.N., P.B., and M.H.A. All authors have read and agreed to the published version of the manuscript.

## INSTITUTIONAL REVIEW BOARD STATEMENT

Human blood samples utilized in the presented analyses were collected in accordance with the principles of the Declaration of Helsinki and consistent with the Good Clinical Practice guidelines of the International Conference on Harmonization. The protocol for the ancillary study (study ID number 195) under the TrialNet’s Natural History Study of the Development of Type 1 Diabetes (Pathway to Prevention Study) TN-01 and was approved by TrialNet’s IRB. Blood samples from healthy subjects were collected as part of another study approved by the IRB of the University of Miami (study number 11995-115).

## INFORMED CONSENT STATEMENT

Informed consent was obtained from all subjects from whom samples were collected according to TrialNet TN-01 and 11995-115 study protocols.

## DATA AVAILABILITY STATEMENT

The current study was conducted on parallel multi-omics datasets previously deposited in the following publicly accessible repositories: the NIH Common Fund’s National Metabolomics Data Repository (NMDR; www.metabolomicsworkbench.org) [106], accession #s ST001690 (doi:10.21228/M8B123) for the metabolomics dataset and ST001642 (doi:10.21228/M8ZX18) for the lipidomics dataset; the PRIDE database of ProteomeXchange (https://www.ebi.ac.uk/pride/), accession # PXD023541 for the proteomics dataset; and the Harvard Dataverse repository (doi.org/10.7910/DVN/A2OU24) for the transcriptomics dataset. Augmented quadra-omics datasets and the code for the data imputation and amplification algorithms can be freely accessed and downloaded in R, Python, and MATLAB scripts by clicking here (https://miamiedu-my.sharepoint.com/:f:/g/personal/mabdulreda_miami_edu/EnQ85UNQRL9Cn4Ee4u_tJHsBcFW_pFOx-Uv1cd2yBsEh1A?e=w12qwg).

## ACKNOWLEDGEMENTS

The authors would like to acknowledge first and foremost the donors of the blood samples whose generous donation made these studies possible. The authors thank the TrialNet clinical center team members (past and present) at the University of Miami for their help with sample collection, and the TrialNet’s IRB and Coordinating Center for clearing the ancillary study that enabled access to the samples. The authors also acknowledge the principal investigator and research team members on the study that allowed access to the samples from healthy subjects. The authors are also grateful to Professor Marian Rewers at the Barbara Davis Center for Childhood Diabetes of the University of Colorado for facilitating access to proteomics data from the DAISY cohort. The authors would like to further acknowledge colleagues from the Duke Molecular Physiology Institute at Duke University Medical Center, the Biological Sciences Division at Pacific Northwest National Laboratory, the Miami Integrative Metabolomics Research Center at the 651 University of Miami, and Ocean Ridge Biosciences for previous assistance in generating the multi-omics datasets upon which the current work was conducted.

