## Supplemental Figures for "A Composite Biomarker Signature of Type 1 Diabetes Risk Identified via Augmentation of Parallel Multi-Omics Data from a Small Cohort"

Presented in these supplementary materials are Table S1, Table S2, Figure S1, Figure S2**,** and Figure S3.

**Table S1:** List of all proteins contributing to the various degrees of overlap in the canonical pathways identified in the original (O-HR) and augmented (A3-HR) proteomics data from the small cohort and previously performed proteomics on the C3 (T2) and C4 (I2) subject groups of the TEDDY cohort [1]. Proteins common across all four datasets, which contributed to the overlap, were highlighted in bold and with an underline. Proteins are listed with their associated gene names.

| Pathway | Original | Amplified | TEDDY-C3 | TEDDY-C4 |
|---|---|---|---|---|
| Actin Cytoskeleton Signaling | None | ACTA1, **ACTB**, **ACTN1**, ARPC5, **CD14**, CRK, **CSK**, EGF, FGD3, GIT1, GNA13, GRB2, **KNG1**, **MYH14**, MYL1, **MYLK**, PAK2, PDGFA, PDGFD, PPP1R12A, RAP2B, RHOA, ROCK1, ROCK2, RRAS, **TLN1**, TLN2, WASF2 | ACTA2, **ACTB**, **ACTN1**, ACTN4, ACTR2, ACTR3, ARPC1B, ARPC2, ARPC3, ARPC4, **CD14**, CFL1, **CSK**, CYFIP1, DIAPH1, EZR, F2, FLNA, FN1, GSN, IQGAP1, ITGA2, ITGA2B, ITGA5, ITGA6, ITGB1, ITGB2, ITGB3, **KNG1**, LBP, MSN, MYH10, MYH11, **MYH14**, MYH9, MYL12A, MYL6, MYL6B, MYL9, **MYLK**, PFN1, RAP1A, **TLN1**, TMSB10/TMSB4X, VCL | ACTA2, **ACTB**, **ACTN1,** ACTN4, ACTR2, ACTR3, ARPC1B, ARPC2, ARPC3, ARPC4, **CD14**, CFL1, **CSK**, CYFIP1, DIAPH1, EZR, F2, FLNA, FN1, GSN, IQGAP1, ITGA2, ITGA2B, ITGA5, ITGA6, ITGB1, ITGB2, ITGB3, **KNG1**, LBP, MSN, MYH10, MYH11, **MYH14**, MYH9, MYL12A, MYL6, MYL6B, MYL9, **MYLK**, PFN1, RAP1A, **TLN1**, TMSB10/TMSB4X, VCL |
| Acute Phase Response Signaling | None | AKT1, **APOA1**, **C4BPA**, **C4BPB**, GRB2, **HNRNPK**, **HRG**, IL18, **ITIH2**, **KLKB1**, MBL2, **OSMR**, RAP2B, RIPK1, RRAS, **SAA1**, **SAA2**, TRAF6 | A2M, AGT, AHSG, ALB, AMBP, APCS, **APOA1**, APOA2, APOH, C1QA, C1QB, C1QC, C1R, C1S, C2, C3, C4A/C4B, **C4BPA**, **C4BPB**, C5, C9, CFB, CP, CRP, F2, F8, FGA, FGB, FGG, FN1, **HNRNPK**, HPX, **HRG**, IL1RAP, IL6R, IL6ST, **ITIH2**, ITIH3, ITIH4, **KLKB1**, LBP, MBL2, **OSMR**, PLG, RAP1A, RBP4, **SAA1**, **SAA2**, SAA4, SERPINA1, SERPINA3, SERPIND1, SERPINE1, SERPINF1, SERPINF2, SERPING1, TF, TTR, VWF | A2M, AGT, AHSG, ALB, AMBP, APCS, **APOA1**, APOA2, APOH, C1QA, C1QB, C1QC, C1R, C1S, C2, C3, C4A/C4B, **C4BPA**, **C4BPB**, C5, C9, CFB, CP, CRP, F2, F8, FGA, FGB, FGG, FN1, **HNRNPK**, HPX, **HRG**, IL1RAP, IL6R, IL6ST, **ITIH2**, ITIH3, ITIH4, **KLKB1**, LBP, **OSMR**, PLG, RAP1A, RBP4, **SAA1**, **SAA2**, SAA4, SERPINA1, SERPINA3, SERPIND1, SERPINE1, SERPINF1, SERPINF2, SERPING1, TF, TTR, VWF |
| Agranulocyte Adhesion and Diapedesis | AOC3, CCL14, ICAM2, MMP14, MMP19, MMP2, MMP9, PODXL, SELL, **VCAM1** | ACTA1, **ACTB**, CCL16, **CCL5**, CD99, CXCL12, IL18, MMP14, MMP9, **MYH14**, MYL1, **VCAM1** | ACTA2, **ACTB**, AOC3, C5, CCL14, **CCL5**, CDH5, EZR, FN1, GNAI2, ICAM1, ICAM2, ITGA5, ITGB1, ITGB2, MADCAM1, MMP19, MMP2, MSN, MYH10, MYH11, **MYH14**, MYH9, MYL6, MYL6B, MYL9, PECAM1, PF4, PODXL, PPBP, SELE, SELL, SELP, **VCAM1** | ACTA2, **ACTB**, AOC3, C5, CCL14, **CCL5**, CDH5, EZR, FN1, GNAI2, ICAM1, ICAM2, ITGA5, ITGB1, ITGB2, MADCAM1, MMP19, MMP2, MSN, MYH10, MYH11, **MYH14**, MYH9, MYL6, MYL6B, MYL9, PECAM1, PF4, PODXL, PPBP, SELE, SELL, SELP, **VCAM1** |
| Complement System | None | **C4BPA**, **C4BPB**, CD46, **CD59**, **CR2**, **MASP1**, **MASP2**, MBL2 | C1QA, C1QB, C1QC, C1R, C1S, C2, C3, C4A/C4B, **C4BPA**, **C4BPB**, C5, C6, C7, C8A, C8B, C8G, C9, CD55, **CD59**, CFB, CFD, CFH, CFI, CR1, **CR2**, ITGB2, **MASP1**, **MASP2**, MBL2, SERPING1 | C1QA, C1QB, C1QC, C1R, C1S, C2, C3, C4A/C4B, **C4BPA**, **C4BPB**, C5, C6, C7, C8A, C8B, C8G, C9, CD55, **CD59**, CFB, CFD, CFH, CFI, CR1, **CR2**, ITGB2, **MASP1**, **MASP2**, SERPING1 |
| Fcγ Receptor-mediated Phagocytosis in Macrophages and Monocytes | None | ACTA1, **ACTB**, AKT1, ARPC5, CRK, FYB1, LCP2, LYN, NCK1, NCK2, **PLD4**, PTK2B, **SRC**, **TLN1**, TLN2, VASP | ACTA2, **ACTB**, ACTR2, ACTR3, ARPC1B, ARPC2, ARPC3, ARPC4, EZR, FCGR2A, FCGR3A/FCGR3B, GPLD1, **PLD4,** RAB11A, **SRC**, **TLN1** | ACTA2, **ACTB**, ACTR2, ACTR3, ARPC1B, ARPC2, ARPC3, ARPC4, EZR, FCGR2A, FCGR3A/FCGR3B, GPLD1, **PLD4**, RAB11A, **SRC**, **TLN1**, VASP |
| FXR/RXR Activation | APOA2, APOE, **CETP**, LIPC, **PLTP** | **A1BG**, AKT1, **APOA1**, **APOA4**, **APOB**, **APOC1**, **APOF**, **APOL1**, **APOM**, **CETP**, **FETUB**, IL18, **KNG1**, **LCAT**, **PLTP**, **PON1**, **PON3**, **SAA1**, **SAA2** | **A1BG**, AGT, AHSG, ALB, AMBP, **APOA1**, APOA2, **APOA4**, **APOB**, **APOC1**, APOC2, APOC3, APOC4, APOD, APOE, **APOF**, APOH, **APOL1**, **APOM**, C3, C4A/C4B, C9, **CETP**, CLU, FBP1, **FETUB**, FGA, FGFR4, GC, HPX, ITIH4, **KNG1**, **LCAT**, LIPC, LPL, PCYOX1, **PLTP**, **PON1**, **PON3**, RBP4, **SAA1**, **SAA2**, SAA4, SERPINA1, SERPINF1, SERPINF2, TF, TTR, VTN | **A1BG**, AGT, AHSG, ALB, AMBP, **APOA1**, APOA2, **APOA4**, **APOB**, **APOC1**, APOC2, APOC3, APOC4, APOD, APOE, **APOF**, APOH, **APOL1**, **APOM**, C3, C4A/C4B, C9, **CETP**, CLU, FBP1, **FETUB**, FGA, FGFR4, GC, HPX, ITIH4, **KNG1**, **LCAT**, LIPC, LPL, PCYOX1, **PLTP**, **PON1**, **PON3**, RBP4, **SAA1**, **SAA2**, SAA4, SERPINA1, SERPINF1, SERPINF2, TF, TTR, VTN |

| Pathway | Original | Amplified | TEDDY-C3 | TEDDY-C4 |
|---|---|---|---|---|
| HIF1α Signaling | **IGF1**, MMP14, MMP19, MMP2, MMP9, RAN | AKT1, CUL2, EGF, ELOB, ELOC, **IGF1**, **IGF2**, LDHC, **MET**, MMP14, MMP9, RAN, RAP2B, RRAS | FLT4, GPI, HK1, HSP90AA1, HSPA1L, HSPA2, HSPA4, HSPA5, HSPA8, **IGF1**, **IGF2**, IL6R, KDR, LDHA, LDHB, **MET**, MMP19, MMP2, PKM, RAP1A, SERPINE1, TF, TGFB1 | FLT4, GPI, HK1, HSP90AA1, HSPA1L, HSPA2, HSPA4, HSPA5, HSPA8, **IGF1**, **IGF2**, IL6R, KDR, LDHA, LDHB, **MET**, MMP19, MMP2, PKM, RAP1A, SERPINE1, TF, TGFB1 |
| ID1 Signaling Pathway | **CCN2**, FGFR2, MMP2, MMP9, TGM2 | AKT1, **CCN2**, EGF, **EGFR**, **FGFR1**, GDF2, **IGF2**, LYN, MMP9, **PTK7**, RAP2B, RHOC, RRAS, **SRC** | APP, ATF6, **CCN2**, **EGFR**, **FGFR1**, FGFR4, FN1, **IGF2**, IL6R, MCAM, MMP2, PARK7, PLXNB1, PLXNB2, **PTK7**, RAP1A, S100A9, **SRC**, TGFB1, TGFBR3 | APP, ATF6, **CCN2**, **EGFR**, **FGFR1**, FGFR4, FN1, **IGF2**, IL6R, MCAM, MMP2, PARK7, PLXNB1, PLXNB2, **PTK7,** RAP1A, S100A9, **SRC**, TGFB1, TGFBR3 |
| IL-12 Signaling and Production in Macrophages | None | AKT1, **APOA1**, **APOA4**, **APOB**, **APOC1**, **APOF**, **APOL1**, **APOM**, IL18, **MST1**, **PON1**, **RAB7A**, TRAF6 | ALB, **APOA1**, APOA2, **APOA4**, **APOB**, **APOC1**, APOC2, APOC3, APOC4, APOD, APOE, **APOF**, **APOL1**, **APOM**, C3, CLU, COL11A2, COL18A1, COL1A1, COL1A2, COL2A1, COL3A1, COL5A3, FCGR2A, FCGR3A/FCGR3B, H3-4, IGHG1, IGHG2, IGHG3, IGHM, IGKC, IL6ST, JCHAIN, LPA, LYZ, **MST1**, PCYOX1, **PON1**, **RAB7A**, RBP4, S100A8, SAA4, SELE, SELP, SERPINA1, TGFB1, THBS1 | ALB, **APOA1**, APOA2, **APOA4**, **APOB**, **APOC1**, APOC2, APOC3, APOC4, APOD, APOE, **APOF**, **APOL1**, **APOM**, C3, CLU, COL11A2, COL18A1, COL1A1, COL1A2, COL2A1, COL3A1, COL5A3, FCGR2A, FCGR3A/FCGR3B, H3-4, IGHG1, IGHG2, IGHG3, IGHM, IGKC, IL6ST, JCHAIN, LPA, LYZ, **MST1**, PCYOX1, **PON1**, **RAB7A**, RBP4, S100A8, SAA4, SELE, SELP, SERPINA1, TGFB1, THBS1 |
| IL-15 Production | CSF1R, FGFR2, PTK2B, ROR1, TYRO3 | **CSK**, **EGFR**, **EPHA4**, **FGFR1**, **KIT**, LYN, **MET**, NTRK3, PTK2B, **PTK7**, RET, ROR1, **SRC**, TEC, **TIE1**, TWF1 | AXL, BTK, CSF1R, CSF2RA, **CSK**, **EGFR**, EPHA1, **EPHA4**, EPHB2, **FGFR1**, FGFR4, FLT4, KDR, **KIT**, MERTK, **MET**, NTRK2, PDGFRB, **PTK7**, **SRC**, TEK, **TIE1**, TYRO3 | AXL, BTK, CSF1R, CSF2RA, **CSK**, **EGFR**, EPHA1, **EPHA4**, EPHB2, **FGFR1**, FGFR4, FLT4, KDR, **KIT**, MERTK, **MET**, NTRK2, PDGFRB, **PTK7**, **SRC**, TEK, **TIE1**, TYRO3 |
| IL-8 Signaling | GNAZ, MMP2, MMP9, **PLD4**, PTK2B, **VCAM1** | AKT1, **CR2**, **EGFR**, GNA13, GNAZ, **LASP1**, MMP9, PAK2, **PLD4**, PTK2B, RAP2B, RHOA, RHOC, ROCK1, ROCK2, RRAS, **SRC**, TRAF6, VASP, **VCAM1** | CDH1, **CR2**, **EGFR**, FLT4, GNAI2, GPLD1, ICAM1, IQGAP1, ITGB2, ITGB3, KDR, **LASP1**, MMP2, MPO, MYL9, **PLD4**, RAP1A, **SRC**, TEK, **VCAM1** | CDH1, **CR2**, **EGFR**, FLT4, GNAI2, GPLD1, ICAM1, IQGAP1, ITGB2, ITGB3, KDR, **LASP1**, MMP2, MPO, MYL9, **PLD4**, RAP1A, **SRC**, TEK, VASP, **VCAM1** |
| Leukocyte Extravasation Signaling | CD44, **ICAM3**, MMP14, MMP19, MMP2, MMP9, PTK2B, TIMP2, **VCAM1** | ACTA1, **ACTB**, **ACTN1**, CD99, CRK, CXCL12, **ICAM3**, MMP14, MMP9, PTK2B, RHOA, ROCK1, ROCK2, **SRC**, TEC, VASP, **VCAM1** | ACTA2, **ACTB**, **ACTN1**, ACTN4, ARHGAP1, BTK, CD44, CDH5, CTTN, EZR, GNAI2, ICAM1, **ICAM3**, ITGA2, ITGB1, ITGB2, MMP19, MMP2, MSN, MYL6, PECAM1, RAP1A, **SRC**, TIMP1, TIMP2, **VCAM1**, VCL | ACTA2, **ACTB**, **ACTN1**, ACTN4, ARHGAP1, BTK, CD44, CDH5, CTTN, EZR, GNAI2, ICAM1, **ICAM3**, ITGA2, ITGB1, ITGB2, MMP19, MMP2, MSN, MYL6, PECAM1, RAP1A, **SRC**, TIMP1, TIMP2, VASP, **VCAM1**, VCL |
| LXR/RXR Activation | APOA2, APOE, **CETP**, LDLR, MMP9, **PLTP** | **A1BG**, **APOA1**, **APOA4**, **APOA5**, **APOB**, **APOC1**, **APOF**, **APOL1**, **APOM**, **CD14**, CD36, **CETP**, IL18, **KNG1**, **LCAT**, MMP9, **PLTP**, **PON1**, **PON3**, **SAA1**, **SAA2** | **A1BG**, AGT, AHSG, ALB, AMBP, **APOA1**, APOA2, **APOA4**, **APOA5**, **APOB**, **APOC1**, APOC2, APOC3, APOC4, APOD, APOE, **APOF**, APOH, **APOL1**, **APOM**, C3, C4A/C4B, C9, **CD14**, **CETP**, CLU, FGA, GC, HPX, IL1R2, IL1RAP, ITIH4, **KNG1**, LBP, **LCAT**, LDLR, LPA, LPL, LYZ, PCYOX1, **PLTP**, **PON1**, **PON3**, RBP4, S100A8, **SAA1**, **SAA2**, SAA4, SERPINA1, SERPINF1, SERPINF2, TF, TTR, VTN | **A1BG**, AGT, AHSG, ALB, AMBP, **APOA1**, APOA2, **APOA4**, **APOA5**, **APOB**, **APOC1**, APOC2, APOC3, APOC4, APOD, APOE, **APOF**, APOH, **APOL1**, **APOM**, C3, C4A/C4B, C9, **CD14**, **CETP**, CLU, FGA, GC, HPX, IL1R2, IL1RAP, ITIH4, **KNG1**, LBP, **LCAT**, LDLR, LPA, LPL, LYZ, PCYOX1, **PLTP**, **PON1**, **PON3**, RBP4, S100A8, **SAA1**, **SAA2**, SAA4, SERPINA1, SERPINF1, SERPINF2, TF, TTR, VTN |

| Pathway | Original | Amplified | TEDDY-C3 | TEDDY-C4 |
|---|---|---|---|---|
| Maturity Onset Diabetes of Young (MODY) Signaling | None | **APOA1**, **APOA4**, **APOA5**, **APOB**, **APOC1**, **APOF**, **APOL1**, **APOM** | ADIPOQ, ALDOB, **APOA1**, APOA2, **APOA4**, **APOA5**, **APOB**, **APOC1**, APOC2, APOC3, APOC4, APOD, APOE, **APOF**, APOH, **APOL1**, **APOM**, CACNA2D1, FABP1, GAPDH | ADIPOQ, ALDOB, **APOA1**, APOA2, **APOA4**, **APOA5**, **APOB**, **APOC1**, APOC2, APOC3, APOC4, APOD, APOE, **APOF**, APOH, **APOL1**, **APOM**, CACNA2D1, FABP1, GAPDH |
| MSP-RON Signaling Pathway | None | ACTA1, **ACTB**, **CSF1**, **KLK6**, **KLKB1**, **MST1** | ACTA2, **ACTB**, **CSF1**, F11, F12, ITGB2, **KLK6**, **KLKB1**, **MST1** | ACTA2, **ACTB**, **CSF1**, F11, F12, ITGB2, **KLK6**, **KLKB1**, **MST1** |
| Natural Killer Cell Signaling | None | AKT1, GRB2, IL18, LCP2, NCK1, PAK2, PTK2B, RAP2B, ROCK1, RRAS, TRAF6 | B2M, CFL1, COL11A2, COL18A1, COL1A1, COL1A2, COL2A1, COL3A1, COL5A3, FCGR2A, FCGR3A/FCGR3B, HLA-B, HSPA1L, HSPA2, HSPA4, HSPA5, HSPA8, ITGB1, PTPN6, PVR, RAP1A | B2M, CFL1, COL11A2, COL18A1, COL1A1, COL1A2, COL2A1, COL3A1, COL5A3, FCGR2A, FCGR3A/FCGR3B, HLA-B, HSPA1L, HSPA2, HSPA4, HSPA5, HSPA8, ITGB1, PTPN6, PVR, RAP1A |
| NRF2-mediated Oxidative Stress Response | None | ACTA1, **ACTB**, AKT1, **CBR1**, **ERP29**, GSTM3, **GSTO1**, **GSTP1**, **NQO2**, **PRDX1**, RAP2B, RRAS, **SOD1**, UBE2K | ACTA2, **ACTB**, CAT, **CBR1**, CCT7, **ERP29**, GSR, **GSTO1**, **GSTP1**, HSP90AA1, HSP90AB1, HSP90B1, **NQO2**, PPIB, **PRDX1**, RAP1A, **SOD1**, SOD3, STIP1, TXN, VCP | ACTA2, **ACTB**, CAT, **CBR1**, CCT7, **ERP29**, GSR, **GSTO1**, **GSTP1**, HSP90AA1, HSP90AB1, HSP90B1, **NQO2**, PPIB, **PRDX1**, RAP1A, **SOD1**, SOD3, STIP1, TXN, VCP |
| Phagosome Formation | None | **ADGRE5**, AKT1, AP1B1, AP1G1, AP1M1, ARPC5, **CD14**, CD36, CLIP1, **CR2**, GRB2, HTR5A, **IGHG2**, IGHG4, **IGHM**, IGLC3, **LCAT**, LYN, **MARCO**, **MYH14**, MYL1, **MYLK**, PAK2, **PLD4**, PTK2B, RAP2B, RHOA, ROCK1, ROCK2, RRAS, **SRC**, **TIMD4**, **TLN1**, TLN2, WASF2 | ACTR2, ACTR3, **ADGRE5**, ADGRF5, ADGRG2, ADGRG6, ADGRL4, ARPC1B, ARPC2, ARPC3, ARPC4, C3, **CD14**, CD209, CFL1, COLEC12, CR1, **CR2**, DIAPH1, FCGR2A, FCGR3A/FCGR3B, FN1, GPLD1, IGHA1, IGHG1, **IGHG2**, IGHG3, **IGHM**, IGKC, ITGA2, ITGA2B, ITGA5, ITGA6, ITGB1, ITGB2, ITGB3, JCHAIN, LBP, **LCAT**, **MARCO**, MRC1, MRC2, MYH10, MYH11, **MYH14**, MYH9, MYL12A, MYL6, MYL6B, MYL9, **MYLK**, PIP4K2A, PLA2G7, **PLD4**, PRDX6, RAP1A, SCARA5, **SRC**, **TIMD4**, **TLN1**, VTN | ACTR2, ACTR3, **ADGRE5**, ADGRF5, ADGRG2, ADGRG6, ADGRL4, ARPC1B, ARPC2, ARPC3, ARPC4, C3, **CD14**, CD209, CFL1, COLEC12, CR1, **CR2**, DIAPH1, FCGR2A, FCGR3A/FCGR3B, FN1, GPLD1, IGHA1, IGHG1, **IGHG2**, IGHG3, **IGHM**, IGKC, ITGA2, ITGA2B, ITGA5, ITGA6, ITGB1, ITGB2, ITGB3, JCHAIN, LBP, **LCAT**, **MARCO**, MRC1, MRC2, MYH10, MYH11, **MYH14**, MYH9, MYL12A, MYL6, MYL6B, MYL9, **MYLK**, PIP4K2A, PLA2G7, **PLD4**, PRDX6, RAP1A, SCARA5, **SRC**, **TIMD4**, **TLN1**, VTN |
| Phagosome Maturation | NSF, **PRDX1**, **PRDX2**, PRDX6, RILP | DCTN4, **LAMP1**, NSF, **PRDX1**, **PRDX2**, RAB5A, **RAB7A**, YKT6 | ATP6V1A, B2M, CALR, CTSA, CTSB, CTSC, CTSD, CTSF, CTSH, CTSL, CTSS, CTSZ, HLA-B, **LAMP1**, LAMP2, MPO, **PRDX1**, **PRDX2**, PRDX5, PRDX6, **RAB7A**, TUBA1A, TUBA1B, TUBA4A, TUBB, TUBB1, TUBB2A | ATP6V1A, B2M, CALR, CTSA, CTSB, CTSC, CTSD, CTSF, CTSH, CTSL, CTSS, CTSZ, HLA-B, **LAMP1**, LAMP2, MPO, **PRDX1**, **PRDX2**, PRDX5, PRDX6, **RAB7A**, TUBA1A, TUBA1B, TUBA4A, TUBB, TUBB1, TUBB2A |
| PPARα/RXRα Activation | None | **APOA1**, **CAND1**, CD36, CKAP5, GRB2, PLCB3, RAP2B, RRAS, TRAF6 | ADIPOQ, **APOA1**, APOA2, **CAND1**, GHR, HSP90AA1, HSP90AB1, HSP90B1, IL1R2, IL1RAP, LPL, PDIA3, PRKAR1A, RAP1A, TGFB1, TGFBR3 | ADIPOQ, **APOA1**, APOA2, **CAND1**, GHR, HSP90AA1, HSP90AB1, HSP90B1, IL1R2, IL1RAP, LPL, PDIA3, PRKAR1A, RAP1A, TGFB1, TGFBR3 |
| Production of Nitric Oxide and Reactive Oxygen Species in Macrophages | None | AKT1, **APOA1**, **APOA4**, **APOB**, **APOC1**, **APOF**, **APOL1**, **APOM**, **PON1**, PPP1R12A, RHOA, RHOC | ALB, **APOA1**, APOA2, **APOA4**, **APOB**, **APOC1**, APOC2, APOC3, APOC4, APOD, APOE, **APOF**, **APOL1**, **APOM**, CAT, CLU, LPA, LYZ, MPO, PCYOX1, **PON1**, PPP1CA, PPP2R1A, PTPN6, RAP1A, RBP4, S100A8, SAA4, SERPINA1, SIRPA | ALB, **APOA1**, APOA2, **APOA4**, **APOB**, **APOC1**, APOC2, APOC3, APOC4, APOD, APOE, **APOF**, **APOL1**, **APOM**, CAT, CLU, LPA, LYZ, MPO, PCYOX1, **PON1**, PPP1CA, PPP2R1A, PTPN6, RAP1A, RBP4, S100A8, SAA4, SERPINA1, SIRPA |

| Pathway | Original | Amplified | TEDDY-C3 | TEDDY-C4 |
|---|---|---|---|---|
| RHOA Signaling | None | ACTA1, **ACTB**, ARPC5, GNA13, **IGF1**, KTN1, MYL1, **MYLK**, PPP1R12A, PTK2B, RHOA, ROCK1, ROCK2, SEPTIN5, **SEPTIN6**, SEPTIN9 | ACTA2, **ACTB**, ACTR2, ACTR3, ARHGAP1, ARPC1B, ARPC2, ARPC3, ARPC4, CFL1, EPHA1, EZR, **IGF1**, MSN, MYL12A, MYL6, MYL6B, MYL9, **MYLK**, NRP2, PFN1, PIP4K2A, SEMA3F, SEPTIN11, **SEPTIN6**, SEPTIN7 | ACTA2, **ACTB**, ACTR2, ACTR3, ARHGAP1, ARPC1B, ARPC2, ARPC3, ARPC4, CFL1, EPHA1, EZR, **IGF1**, MSN, MYL12A, MYL6, MYL6B, MYL9, **MYLK**, NRP2, PFN1, PIP4K2A, SEMA3F, SEPTIN11, **SEPTIN6**, SEPTIN7 |
| RHOGDI Signaling | None | ACTA1, **ACTB**, ARPC5, GNA13, GNAZ, **MYH14**, MYL1, PAK2, PPP1R12A, RHOA, RHOC, ROCK1, ROCK2, **SRC**, WASF2 | ACTA2, **ACTB**, ACTR2, ACTR3, ARHGAP1, ARHGDIA, ARHGDIB, ARPC1B, ARPC2, ARPC3, ARPC4, CD44, CDH1, CDH11, CDH13, CDH15, CDH17, CDH2, CDH5, CDH6, CFL1, EZR, GNAI2, ITGA2, ITGA2B, ITGA5, ITGA6, ITGB1, ITGB2, ITGB3, MSN, MYH10, MYH11, **MYH14**, MYH9, MYL12A, MYL6, MYL6B, MYL9, PIP4K2A, **SRC** | ACTA2, **ACTB**, ACTR2, ACTR3, ARHGAP1, ARHGDIA, ARHGDIB, ARPC1B, ARPC2, ARPC3, ARPC4, CD44, CDH1, CDH11, CDH13, CDH15, CDH17, CDH2, CDH5, CDH6, CFL1, EZR, GNAI2, ITGA2, ITGA2B, ITGA5, ITGA6, ITGB1, ITGB2, ITGB3, MSN, MYH10, MYH11, **MYH14**, MYH9, MYL12A, MYL6, MYL6B, MYL9, PIP4K2A, **SRC** |
| Role of Macrophages, Fibroblasts and Endothelial Cells in Rheumatoid Arthritis | None | AKT1, CALM1, **CCL5**, **CSF1**, CXCL12, **IGHG2**, IGHG4, IGLC3, IL18, **LRP1**, LRP6, PDGFA, PDGFD, PLCB3, RAP2B, RHOA, RIPK1, ROCK1, ROCK2, RRAS, **SRC**, TRAF6, **VCAM1** | C5, **CCL5**, **CSF1**, DKK3, FCGR3A/FCGR3B, FN1, ICAM1, IGHG1, **IGHG2**, IGHG3, IGHM, IGKC, IL1R2, IL1RAP, IL6R, IL6ST, JCHAIN, **LRP1**, MIF, PDIA3, PRSS1, RAP1A, SELE, **SRC**, TGFB1, **VCAM1** | C5, **CCL5**, **CSF1**, DKK3, FCGR3A/FCGR3B, FN1, ICAM1, IGHG1, **IGHG2**, IGHG3, IGHM, IGKC, IL1R2, IL1RAP, IL6R, IL6ST, JCHAIN, **LRP1**, MIF, PDIA3, PRSS1, RAP1A, SELE, **SRC**, TGFB1, **VCAM1** |
| Role of PKR in Interferon Induction and Antiviral Response | None | BID, CASP3, FADD, FAS, IL18, **MARCO**, PDGFA, PDGFD, TRAF6 | COLEC12, HSP90AA1, HSP90AB1, HSP90B1, HSPA1L, HSPA2, HSPA4, HSPA5, HSPA8, **MARCO**, PDGFRB, SCARA5 | COLEC12, HSP90AA1, HSP90AB1, HSP90B1, HSPA1L, HSPA2, HSPA4, HSPA5, HSPA8, **MARCO**, PDGFRB, SCARA5 |
| Signaling by Rho Family GTPases | None | ACTA1, **ACTB**, ARPC5, CLIP1, GFAP, GNA13, GNAZ, MYL1, **MYLK**, PAK2, PPP1R12A, PTK2B, RHOA, RHOC, ROCK1, ROCK2, SEPTIN5, **SEPTIN6**, SEPTIN9 | ACTA2, **ACTB**, ACTR2, ACTR3, ARPC1B, ARPC2, ARPC3, ARPC4, CDH1, CDH11, CDH13, CDH15, CDH17, CDH2, CDH5, CDH6, CFL1, CYFIP1, EZR, GNAI2, IQGAP1, ITGA2, ITGA2B, ITGA5, ITGA6, ITGB1, ITGB2, ITGB3, MSN, MYL12A, MYL6, MYL6B, MYL9, **MYLK**, PIP4K2A, SEPTIN11, **SEPTIN6**, SEPTIN7 | ACTA2, **ACTB**, ACTR2, ACTR3, ARPC1B, ARPC2, ARPC3, ARPC4, CDH1, CDH11, CDH13, CDH15, CDH17, CDH2, CDH5, CDH6, CFL1, CYFIP1, EZR, GNAI2, IQGAP1, ITGA2, ITGA2B, ITGA5, ITGA6, ITGB1, ITGB2, ITGB3, MSN, MYL12A, MYL6, MYL6B, MYL9, **MYLK**, PIP4K2A, SEPTIN11, **SEPTIN6**, SEPTIN7 |
| STAT3 Pathway | None | EGF, **EGFR**, **FGFR1**, **IGF1**, NTRK3, RAP2B, RRAS, **SRC** | **EGFR**, **FGFR1**, FGFR4, FLT4, GHR, **IGF1**, IGF2R, IL13RA1, IL1R2, IL6R, IL6ST, KDR, NTRK2, PDGFRB, PTPN6, RAP1A, **SRC**, TGFB1, TGFBR3 | **EGFR**, **FGFR1**, FGFR4, FLT4, GHR, **IGF1**, IGF2R, IL13RA1, IL1R2, IL6R, IL6ST, KDR, NTRK2, PDGFRB, PTPN6, RAP1A, **SRC**, TGFB1, TGFBR3 |
| Tumor Microenvironment Pathway | CD44, **CSPG4**, **IGF1**, MMP14, MMP19, MMP2, MMP9 | AKT1, **CSF1**, **CSPG4**, CXCL12, EGF, FAS, **IGF1**, **IGF2**, MMP14, MMP9, PDGFA, PDGFD, RAP2B, RRAS | CD44, COL1A1, COL1A2, COL3A1, **CSF1**, **CSPG4**, FN1, HLA-B, ICAM1, **IGF1**, **IGF2**, IL6R, ITGA5, ITGB3, LEPR, MMP19, MMP2, RAP1A, SPP1, TGFB1, TNC | CD44, COL1A1, COL1A2, COL3A1, **CSF1**, **CSPG4**, FN1, HLA-B, ICAM1, **IGF1**, **IGF2**, IL6R, ITGA5, ITGB3, LEPR, MMP19, MMP2, RAP1A, SPP1, TGFB1, TNC |
| VEGF Signaling | None | ACTA1, **ACTB**, **ACTN1**, AKT1, GRB2, PTK2B, RAP2B, ROCK1, ROCK2, RRAS, **SRC** | ACTA2, **ACTB**, **ACTN1**, ACTN4, FLT4, KDR, PTPN6, RAP1A, **SRC**, VCL, YWHAE | ACTA2, **ACTB**, **ACTN1**, ACTN4, FLT4, KDR, PTPN6, RAP1A, **SRC**, VCL, YWHAE |

End of table

**Table S2:** List of all peptides identified with post-translational modifications (including glycation) as part of the proteomics analysis performed in the plasma samples of high-risk subjects in the small cohort of samples utilized in the quadra-omics studies and the data augmentation approach [2, 3]. Symbols for post-translational modifications are as follows: hashtag (#) for glycation/glycozylation and asterisk (*) for osidation. Dot (.) indicates enzymatic cleavage.

| Accession | Protein names | Gene | Peptide |
|---|---|---|---|
| P04217 | Alpha-1B-glycoprotein (Alpha-1-B glycoprotein) | A1BG | K.SLPAPWLSMAPVSWITPGLK#TTA.V |
| P04217 | Alpha-1B-glycoprotein (Alpha-1-B glycoprotein) | A1BG | R.LHDNQNGWSGDSAPVELILSDETLPAPEFSPEPESGR#ALR#LR#C*LAPLEG.A |
| P04217 | Alpha-1B-glycoprotein (Alpha-1-B glycoprotein) | A1BG | R.YRLHDNQNGWSGDSAPVELILSDETLPAPEFSPEPESGR#ALR#LR#CLAPL.E |
| P04217 | Alpha-1B-glycoprotein (Alpha-1-B glycoprotein) | A1BG | Y.R#LHDNQNGW*SGDSAPVELILSDETLPAPEFSPEPESGR.A |
| P63261 | Actin, cytoplasmic 2 (EC 3.6.4.-) (Gamma-actin) [Cleaved into: Actin, cytoplasmic 2, N-terminally processed] | ACTG1 ACTG | C.DVDIRK#DLYANTVLSGGTTMYPGIADR.M |
| P43652 | Afamin (Alpha-albumin) (Alpha-Alb) | AFM ALB2 ALBA | C.CK#AESPEVCFNEESPKIGN.- |
| P43652 | Afamin (Alpha-albumin) (Alpha-Alb) | AFM ALB2 ALBA | C.MADK#TLPECSKLPNNVLQEK.I |
| P43652 | Afamin (Alpha-albumin) (Alpha-Alb) | AFM ALB2 ALBA | K.ELISLVEDVSSNYDGC*C*EGDVVQC*IR#.D |
| P43652 | Afamin (Alpha-albumin) (Alpha-Alb) | AFM ALB2 ALBA | K.LKHELTDEELQSLFTNFANVVDK#.C |
| P43652 | Afamin (Alpha-albumin) (Alpha-Alb) | AFM ALB2 ALBA | K.LKHELTDEELQSLFTNFANVVDK#CC*KAESPEVC*FNE.E |
| P02765 | Alpha-2-HS-glycoprotein (Alpha-2-Z-globulin) (Ba-alpha-2-glycoprotein) (Fetuin-A) [Cleaved into: Alpha-2-HS-glycoprotein chain A; Alpha-2-HS-glycoprotein chain B] | AHSG FETUA PRO2743 | G.PGLIY*R#QPNCDDPETEEAALVAIDYINQNLPWGYK.H |
| P02765 | Alpha-2-HS-glycoprotein (Alpha-2-Z-globulin) (Ba-alpha-2-glycoprotein) (Fetuin-A) [Cleaved into: Alpha-2-HS-glycoprotein chain A; Alpha-2-HS-glycoprotein chain B] | AHSG FETUA PRO2743 | H.GPGLIY*R#QPNC*DDPETEEAALVAIDYINQNLPWGYK.H |
| P02765 | Alpha-2-HS-glycoprotein (Alpha-2-Z-globulin) (Ba-alpha-2-glycoprotein) (Fetuin-A) [Cleaved into: Alpha-2-HS-glycoprotein chain A; Alpha-2-HS-glycoprotein chain B] | AHSG FETUA PRO2743 | H.GPGLIY*R#QPNCDDPETEEAALVAIDY*INQNLPW*GYK.H |
| P02765 | Alpha-2-HS-glycoprotein (Alpha-2-Z-globulin) (Ba-alpha-2-glycoprotein) (Fetuin-A) [Cleaved into: Alpha-2-HS-glycoprotein chain A; Alpha-2-HS-glycoprotein chain B] | AHSG FETUA PRO2743 | H.GPGLIY*R#QPNCDDPETEEAALVAIDY*INQNLPWGYK.H |
| P02765 | Alpha-2-HS-glycoprotein (Alpha-2-Z-globulin) (Ba-alpha-2-glycoprotein) (Fetuin-A) [Cleaved into: Alpha-2-HS-glycoprotein chain A; Alpha-2-HS-glycoprotein chain B] | AHSG FETUA PRO2743 | H.GPGLIY*R#QPNCDDPETEEAALVAIDYINQNLPW*GYK.H |
| P02765 | Alpha-2-HS-glycoprotein (Alpha-2-Z-globulin) (Ba-alpha-2-glycoprotein) (Fetuin-A) [Cleaved into: Alpha-2-HS-glycoprotein chain A; Alpha-2-HS-glycoprotein chain B] | AHSG FETUA PRO2743 | H.GPGLIY*R#QPNCDDPETEEAALVAIDYINQNLPWGYK.H |
| P02765 | Alpha-2-HS-glycoprotein (Alpha-2-Z-globulin) (Ba-alpha-2-glycoprotein) (Fetuin-A) [Cleaved into: Alpha-2-HS-glycoprotein chain A; Alpha-2-HS-glycoprotein chain B] | AHSG FETUA PRO2743 | H.GPGLIY*R#QPNCDDPETEEAALVAIDYINQNLPWGYKHTLNQIDEVK.V |
| P02765 | Alpha-2-HS-glycoprotein (Alpha-2-Z-globulin) (Ba-alpha-2-glycoprotein) (Fetuin-A) [Cleaved into: Alpha-2-HS-glycoprotein chain A; Alpha-2-HS-glycoprotein chain B] | AHSG FETUA PRO2743 | H.GPGLIYR#QPNC*DDPETEEAALVAIDYINQNLPW*GYK.H |
| P02765 | Alpha-2-HS-glycoprotein (Alpha-2-Z-globulin) (Ba-alpha-2-glycoprotein) (Fetuin-A) [Cleaved into: Alpha-2-HS-glycoprotein chain A; Alpha-2-HS-glycoprotein chain B] | AHSG FETUA PRO2743 | H.GPGLIYR#QPNC*DDPETEEAALVAIDYINQNLPWGYK.H |
| P02765 | Alpha-2-HS-glycoprotein (Alpha-2-Z-globulin) (Ba-alpha-2-glycoprotein) (Fetuin-A) [Cleaved into: Alpha-2-HS-glycoprotein chain A; Alpha-2-HS-glycoprotein chain B] | AHSG FETUA PRO2743 | K.CNLLAEK#QYGFCK.A |
| P02765 | Alpha-2-HS-glycoprotein (Alpha-2-Z-globulin) (Ba-alpha-2-glycoprotein) (Fetuin-A) [Cleaved into: Alpha-2-HS-glycoprotein chain A; Alpha-2-HS-glycoprotein chain B] | AHSG FETUA PRO2743 | K.EHAVEGDCDFQLLKLDGK#FSVVYAKC*D.S |
| P02765 | Alpha-2-HS-glycoprotein (Alpha-2-Z-globulin) (Ba-alpha-2-glycoprotein) (Fetuin-A) [Cleaved into: Alpha-2-HS-glycoprotein chain A; Alpha-2-HS-glycoprotein chain B] | AHSG FETUA PRO2743 | K.HTLNQIDEVK#VW*PQQPSGE.L |
| P02765 | Alpha-2-HS-glycoprotein (Alpha-2-Z-globulin) (Ba-alpha-2-glycoprotein) (Fetuin-A) [Cleaved into: Alpha-2-HS-glycoprotein chain A; Alpha-2-HS-glycoprotein chain B] | AHSG FETUA PRO2743 | K.HTLNQIDEVK#VW*PQQPSGELFEIEIDTLETTCHVLDPTPVAR.C |
| P02765 | Alpha-2-HS-glycoprotein (Alpha-2-Z-globulin) (Ba-alpha-2-glycoprotein) (Fetuin-A) [Cleaved into: Alpha-2-HS-glycoprotein chain A; Alpha-2-HS-glycoprotein chain B] | AHSG FETUA PRO2743 | K.HTLNQIDEVK#VWPQQPSGELFEIEIDTLETTCHVLDPTPVAR.C |
| P02765 | Alpha-2-HS-glycoprotein (Alpha-2-Z-globulin) (Ba-alpha-2-glycoprotein) (Fetuin-A) [Cleaved into: Alpha-2-HS-glycoprotein chain A; Alpha-2-HS-glycoprotein chain B] | AHSG FETUA PRO2743 | P.GLIYR#QPNCDDPETEEAALVAIDYINQNLPWGYK.H |
| P02765 | Alpha-2-HS-glycoprotein (Alpha-2-Z-globulin) (Ba-alpha-2-glycoprotein) (Fetuin-A) [Cleaved into: Alpha-2-HS-glycoprotein chain A; Alpha-2-HS-glycoprotein chain B] | AHSG FETUA PRO2743 | Q.IDEVK#VW*PQQPSGELFEIEIDTLETTCHVLDPTPVAR.C |
| P02765 | Alpha-2-HS-glycoprotein (Alpha-2-Z-globulin) (Ba-alpha-2-glycoprotein) (Fetuin-A) [Cleaved into: Alpha-2-HS-glycoprotein chain A; Alpha-2-HS-glycoprotein chain B] | AHSG FETUA PRO2743 | R.QLK#EHAVEGDCDFQLLK.L |
| P02765 | Alpha-2-HS-glycoprotein (Alpha-2-Z-globulin) (Ba-alpha-2-glycoprotein) (Fetuin-A) [Cleaved into: Alpha-2-HS-glycoprotein chain A; Alpha-2-HS-glycoprotein chain B] | AHSG FETUA PRO2743 | R.QPNCDDPETEEAALVAIDY*INQNLPW*GY*K#.H |

| Accession | Protein names | Gene | Peptide |
|---|---|---|---|
| P02765 | Alpha-2-HS-glycoprotein (Alpha-2-Z-globulin) (Ba-alpha-2-glycoprotein) (Fetuin-A) [Cleaved into: Alpha-2-HS-glycoprotein chain A; Alpha-2-HS-glycoprotein chain B] | AHSG FETUA PRO2743 | S.APHGPGLIYRQPNCDDPETEEAALVAIDY*INQNLPW*GY*K#.H |
| P02760 | Protein AMBP (Protein HC) [Cleaved into: Alpha-1-microglobulin (EC 1.6.2.-) (Alpha-1 microglycoprotein) (Complex-forming glycoprotein heterogeneous in charge); Inter-alpha-trypsin inhibitor light chain (ITI-LC) (Bikunin) (EDC1) (HI-30) (Uronic-acid-rich p | AMBP HCP ITIL | K.FYSEKEC*R#EY*C*GVPGDGDEELLR.F |
| P02760 | Protein AMBP (Protein HC) [Cleaved into: Alpha-1-microglobulin (EC 1.6.2.-) (Alpha-1 microglycoprotein) (Complex-forming glycoprotein heterogeneous in charge); Inter-alpha-trypsin inhibitor light chain (ITI-LC) (Bikunin) (EDC1) (HI-30) (Uronic-acid-rich p | AMBP HCP ITIL | R.VVAQGVGIPEDSIFTM*ADR#GEC*VPGEQEPEPILIP.R |
| P15144 | Aminopeptidase N (AP-N) (hAPN) (EC 3.4.11.2) (Alanyl aminopeptidase) (Aminopeptidase M) (AP-M) (Microsomal aminopeptidase) (Myeloid plasma membrane glycoprotein CD13) (gp150) (CD antigen CD13) | ANPEP APN CD13 PEPN | S.LVK#DSQYEMDSEFEGELADDLAGFYR.S |
| P04114 | Apolipoprotein B-100 (Apo B-100) [Cleaved into: Apolipoprotein B-48 (Apo B-48)] | APOB | E.SDEETQIK#VNW*EEEAASGLLTSLKDNVPK.A |
| P04114 | Apolipoprotein B-100 (Apo B-100) [Cleaved into: Apolipoprotein B-48 (Apo B-48)] | APOB | E.SDEETQIK#VNWEEEAASGLLTSLK.D |
| P04114 | Apolipoprotein B-100 (Apo B-100) [Cleaved into: Apolipoprotein B-48 (Apo B-48)] | APOB | E.SDEETQIK#VNWEEEAASGLLTSLKDNVPK.A |
| P04114 | Apolipoprotein B-100 (Apo B-100) [Cleaved into: Apolipoprotein B-48 (Apo B-48)] | APOB | K.GISTSAASPAVGTVGMDMDEDDDFSK#W*NFYYSPQSSP.D |
| P04114 | Apolipoprotein B-100 (Apo B-100) [Cleaved into: Apolipoprotein B-48 (Apo B-48)] | APOB | R.DLKVEDIPLAR#IT.L |
| P04114 | Apolipoprotein B-100 (Apo B-100) [Cleaved into: Apolipoprotein B-48 (Apo B-48)] | APOB | R.ESDEETQIK#VNWEEEAASGLLTSLKDNVPK.A |
| P02649 | Apolipoprotein E (Apo-E) | APOE | R.WVQTLSEQVQEELLSSQVTQELRALMDETM*KELK#AY*KSELEEQL.T |
| P02649 | Apolipoprotein E (Apo-E) | APOE | S.GQR#WELALGR#FWDYLR#WVQTLSEQVQEELLSSQVTQELR.A |
| P02749 | Beta-2-glycoprotein 1 (APC inhibitor) (Activated protein C-binding protein) (Anticardiolipin cofactor) (Apolipoprotein H) (Apo-H) (Beta-2-glycoprotein I) (B2GPI) (Beta(2)GPI) | APOH B2G1 | C.TEEGK#W*SPELPVCAPIICPPPSIPTFATLR.V |
| P02749 | Beta-2-glycoprotein 1 (APC inhibitor) (Activated protein C-binding protein) (Anticardiolipin cofactor) (Apolipoprotein H) (Apo-H) (Beta-2-glycoprotein I) (B2GPI) (Beta(2)GPI) | APOH B2G1 | C.TEEGK#WSPELPVCAPIIC*PPPSIPTFATLR.V |
| P02749 | Beta-2-glycoprotein 1 (APC inhibitor) (Activated protein C-binding protein) (Anticardiolipin cofactor) (Apolipoprotein H) (Apo-H) (Beta-2-glycoprotein I) (B2GPI) (Beta(2)GPI) | APOH B2G1 | C.TEEGK#WSPELPVCAPIICPPPSIPTFATLR.V |
| P02749 | Beta-2-glycoprotein 1 (APC inhibitor) (Activated protein C-binding protein) (Anticardiolipin cofactor) (Apolipoprotein H) (Apo-H) (Beta-2-glycoprotein I) (B2GPI) (Beta(2)GPI) | APOH B2G1 | K.CFKEHSSLAFWKTDASDVK#P.C |
| P02749 | Beta-2-glycoprotein 1 (APC inhibitor) (Activated protein C-binding protein) (Anticardiolipin cofactor) (Apolipoprotein H) (Apo-H) (Beta-2-glycoprotein I) (B2GPI) (Beta(2)GPI) | APOH B2G1 | K.DKATFGCHDGYSLDGPEEIECTK#L.G |
| P02749 | Beta-2-glycoprotein 1 (APC inhibitor) (Activated protein C-binding protein) (Anticardiolipin cofactor) (Apolipoprotein H) (Apo-H) (Beta-2-glycoprotein I) (B2GPI) (Beta(2)GPI) | APOH B2G1 | K.EHSSLAFWKTDASDVK#P.C |
| P02749 | Beta-2-glycoprotein 1 (APC inhibitor) (Activated protein C-binding protein) (Anticardiolipin cofactor) (Apolipoprotein H) (Apo-H) (Beta-2-glycoprotein I) (B2GPI) (Beta(2)GPI) | APOH B2G1 | K.TFYEPGEEITY*SC*K#PGY*VSR.G |
| P02749 | Beta-2-glycoprotein 1 (APC inhibitor) (Activated protein C-binding protein) (Anticardiolipin cofactor) (Apolipoprotein H) (Apo-H) (Beta-2-glycoprotein I) (B2GPI) (Beta(2)GPI) | APOH B2G1 | K.TFYEPGEEITYSCK#PGY.V |
| P02749 | Beta-2-glycoprotein 1 (APC inhibitor) (Activated protein C-binding protein) (Anticardiolipin cofactor) (Apolipoprotein H) (Apo-H) (Beta-2-glycoprotein I) (B2GPI) (Beta(2)GPI) | APOH B2G1 | K.TFYEPGEEITYSCK#PGYVSR.G |
| P02749 | Beta-2-glycoprotein 1 (APC inhibitor) (Activated protein C-binding protein) (Anticardiolipin cofactor) (Apolipoprotein H) (Apo-H) (Beta-2-glycoprotein I) (B2GPI) (Beta(2)GPI) | APOH B2G1 | K.TFYEPGEEITYSCKPGYVSR#G.G |
| P02749 | Beta-2-glycoprotein 1 (APC inhibitor) (Activated protein C-binding protein) (Anticardiolipin cofactor) (Apolipoprotein H) (Apo-H) (Beta-2-glycoprotein I) (B2GPI) (Beta(2)GPI) | APOH B2G1 | R.TCPKPDDLPFSTVVPLK#TFY*EPGEEITYSCKPGYVS.R |
| P25311 | Zinc-alpha-2-glycoprotein (Zn-alpha-2-GP) (Zn-alpha-2-glycoprotein) | AZGP1 ZAG ZNGP1 | R.QVEGMEDWKQDSQLQK#.A |

| Accession | Protein names | Gene | Peptide |
|---|---|---|---|
| P09871 | Complement C1s subcomponent (EC 3.4.21.42) (C1 esterase) (Complement component 1 subcomponent s) [Cleaved into: Complement C1s subcomponent heavy chain; Complement C1s subcomponent light chain] | C1S | C.K#EVKVEKPTADAEAYVFTPNMICAGGEK.G |
| P09871 | Complement C1s subcomponent (EC 3.4.21.42) (C1 esterase) (Complement component 1 subcomponent s) [Cleaved into: Complement C1s subcomponent heavy chain; Complement C1s subcomponent light chain] | C1S | C.KEVK#VEKPTADAEAYVFTPNMICAGGEK.G |
| P09871 | Complement C1s subcomponent (EC 3.4.21.42) (C1 esterase) (Complement component 1 subcomponent s) [Cleaved into: Complement C1s subcomponent heavy chain; Complement C1s subcomponent light chain] | C1S | C.KEVKVEK#PTADAEAYVFTPNM*ICAGGEK.G |
| P06681 | Complement C2 (EC 3.4.21.43) (C3/C5 convertase) [Cleaved into: Complement C2b fragment; Complement C2a fragment] | C2 | Q.CR#PNGMW*DGETAVCDNGAGHCPNPGISLGAVR.T |
| P01024 | Complement C3 (C3 and PZP-like alpha-2-macroglobulin domain-containing protein 1) [Cleaved into: Complement C3 beta chain; C3-beta-c (C3bc); Complement C3 alpha chain; C3a anaphylatoxin; Acylation stimulating protein (ASP) (C3adesArg); Complement C3b alph | C3 CPAMD1 | K.DICEEQVNSLPGSITKAGDFLEANY*MNLQR#S.Y |
| P01024 | Complement C3 (C3 and PZP-like alpha-2-macroglobulin domain-containing protein 1) [Cleaved into: Complement C3 beta chain; C3-beta-c (C3bc); Complement C3 alpha chain; C3a anaphylatoxin; Acylation stimulating protein (ASP) (C3adesArg); Complement C3b alph | C3 CPAMD1 | K.DTWVEHWPEEDECQDEENQK#QC*QDLGAFTESMVVFGC*P.N |
| P01024 | Complement C3 (C3 and PZP-like alpha-2-macroglobulin domain-containing protein 1) [Cleaved into: Complement C3 beta chain; C3-beta-c (C3bc); Complement C3 alpha chain; C3a anaphylatoxin; Acylation stimulating protein (ASP) (C3adesArg); Complement C3b alph | C3 CPAMD1 | K.GICVADPFEVTVMQDFFIDLR#LPYSVVR.N |
| P01024 | Complement C3 (C3 and PZP-like alpha-2-macroglobulin domain-containing protein 1) [Cleaved into: Complement C3 beta chain; C3-beta-c (C3bc); Complement C3 alpha chain; C3a anaphylatoxin; Acylation stimulating protein (ASP) (C3adesArg); Complement C3b alph | C3 CPAMD1 | N.R#WEDPGKQLYNVEATSYALLALLQLKDFDFVPPVVR.W |
| P01024 | Complement C3 (C3 and PZP-like alpha-2-macroglobulin domain-containing protein 1) [Cleaved into: Complement C3 beta chain; C3-beta-c (C3bc); Complement C3 alpha chain; C3a anaphylatoxin; Acylation stimulating protein (ASP) (C3adesArg); Complement C3b alph | C3 CPAMD1 | R.LDK#ACEPGVDYVYK.T |
| P01024 | Complement C3 (C3 and PZP-like alpha-2-macroglobulin domain-containing protein 1) [Cleaved into: Complement C3 beta chain; C3-beta-c (C3bc); Complement C3 alpha chain; C3a anaphylatoxin; Acylation stimulating protein (ASP) (C3adesArg); Complement C3b alph | C3 CPAMD1 | R.LESEETMVLEAHDAQGDVPVTVTVHDFPGK#K.L |
| P01024 | Complement C3 (C3 and PZP-like alpha-2-macroglobulin domain-containing protein 1) [Cleaved into: Complement C3 beta chain; C3-beta-c (C3bc); Complement C3 alpha chain; C3a anaphylatoxin; Acylation stimulating protein (ASP) (C3adesArg); Complement C3b alph | C3 CPAMD1 | R.VPVAVQGEDTVQSLTQGDGVAK#LSINTHPSQK#PLSIT.V |
| P01024 | Complement C3 (C3 and PZP-like alpha-2-macroglobulin domain-containing protein 1) [Cleaved into: Complement C3 beta chain; C3-beta-c (C3bc); Complement C3 alpha chain; C3a anaphylatoxin; Acylation stimulating protein (ASP) (C3adesArg); Complement C3b alph | C3 CPAMD1 | R.YRGDQDATMSILDISMMTGFAPDTDDLK#QLANG.V |
| P01024 | Complement C3 (C3 and PZP-like alpha-2-macroglobulin domain-containing protein 1) [Cleaved into: Complement C3 beta chain; C3-beta-c (C3bc); Complement C3 alpha chain; C3a anaphylatoxin; Acylation stimulating protein (ASP) (C3adesArg); Complement C3b alph | C3 CPAMD1 | Y.FK#PGMPFDLMVFVTNPDGSPAYR.V |
| P01024 | Complement C3 (C3 and PZP-like alpha-2-macroglobulin domain-containing protein 1) [Cleaved into: Complement C3 beta chain; C3-beta-c (C3bc); Complement C3 alpha chain; C3a anaphylatoxin; Acylation stimulating protein (ASP) (C3adesArg); Complement C3b alph | C3 CPAMD1 | Y.FK#PGMPFDLMVFVTNPDGSPAYRVPVAVQGEDTVQSLTQGDGVAK.L |
| P01024 | Complement C3 (C3 and PZP-like alpha-2-macroglobulin domain-containing protein 1) [Cleaved into: Complement C3 beta chain; C3-beta-c (C3bc); Complement C3 alpha chain; C3a anaphylatoxin; Acylation stimulating protein (ASP) (C3adesArg); Complement C3b alph | C3 CPAMD1 | Y.R#GDQDATM*SILDISMMTGFAPDTDDLK.Q |
| P01024 | Complement C3 (C3 and PZP-like alpha-2-macroglobulin domain-containing protein 1) [Cleaved into: Complement C3 beta chain; C3-beta-c (C3bc); Complement C3 alpha chain; C3a anaphylatoxin; Acylation stimulating protein (ASP) (C3adesArg); Complement C3b alph | C3 CPAMD1 | Y.R#GDQDATMSILDISMM*TGFAPDTDDLK.Q |
| P01024 | Complement C3 (C3 and PZP-like alpha-2-macroglobulin domain-containing protein 1) [Cleaved into: Complement C3 beta chain; C3-beta-c (C3bc); Complement C3 alpha | C3 CPAMD1 | Y.R#GDQDATMSILDISMMTGFAPDTDDLK.Q |

| Accession | Protein names | Gene | Peptide |
|---|---|---|---|
| | chain; C3a anaphylatoxin; Acylation stimulating protein (ASP) (C3adesArg); Complement C3b alph | | |
| P0C0L4 | Complement C4-A (Acidic complement C4) (C3 and PZP-like alpha-2-macroglobulin domain-containing protein 2) [Cleaved into: Complement C4 beta chain; Complement C4-A alpha chain; C4a anaphylatoxin; C4b-A; C4d-A; Complement C4 gamma chain] | C4A CO4 CPAMD2 | K.VLSLAQEQVGGSPEK#LQETSNWLLSQQQADGSFQDPCPVL.D |
| P0C0L4 | Complement C4-A (Acidic complement C4) (C3 and PZP-like alpha-2-macroglobulin domain-containing protein 2) [Cleaved into: Complement C4 beta chain; Complement C4-A alpha chain; C4a anaphylatoxin; C4b-A; C4d-A; Complement C4 gamma chain] | C4A CO4 CPAMD2 | R.ECVGFEAVQEVPVGLVQPASATLY*DY*Y*NPER#.R |
| P0C0L4 | Complement C4-A (Acidic complement C4) (C3 and PZP-like alpha-2-macroglobulin domain-containing protein 2) [Cleaved into: Complement C4 beta chain; Complement C4-A alpha chain; C4a anaphylatoxin; C4b-A; C4d-A; Complement C4 gamma chain] | C4A CO4 CPAMD2 | R.ECVGFEAVQEVPVGLVQPASATLY*DY*YNPER#.R |
| P0C0L4 | Complement C4-A (Acidic complement C4) (C3 and PZP-like alpha-2-macroglobulin domain-containing protein 2) [Cleaved into: Complement C4 beta chain; Complement C4-A alpha chain; C4a anaphylatoxin; C4b-A; C4d-A; Complement C4 gamma chain] | C4A CO4 CPAMD2 | R.LLATLCSAEVC*QC*AEGK#C*PR.Q |
| P0C0L4 | Complement C4-A (Acidic complement C4) (C3 and PZP-like alpha-2-macroglobulin domain-containing protein 2) [Cleaved into: Complement C4 beta chain; Complement C4-A alpha chain; C4a anaphylatoxin; C4b-A; C4d-A; Complement C4 gamma chain] | C4A CO4 CPAMD2 | R.LLLFSPSVVHLGVPLSVGVQLQDVPR#GQV.V |
| P0C0L4 | Complement C4-A (Acidic complement C4) (C3 and PZP-like alpha-2-macroglobulin domain-containing protein 2) [Cleaved into: Complement C4 beta chain; Complement C4-A alpha chain; C4a anaphylatoxin; C4b-A; C4d-A; Complement C4 gamma chain] | C4A CO4 CPAMD2 | R.VGDTLNLNLR#AVG.S |
| P0C0L4 | Complement C4-A (Acidic complement C4) (C3 and PZP-like alpha-2-macroglobulin domain-containing protein 2) [Cleaved into: Complement C4 beta chain; Complement C4-A alpha chain; C4a anaphylatoxin; C4b-A; C4d-A; Complement C4 gamma chain] | C4A CO4 CPAMD2 | R.VQQPDC*R#EPFLSC*C*QFAESLR.K |
| P0C0L4 | Complement C4-A (Acidic complement C4) (C3 and PZP-like alpha-2-macroglobulin domain-containing protein 2) [Cleaved into: Complement C4 beta chain; Complement C4-A alpha chain; C4a anaphylatoxin; C4b-A; C4d-A; Complement C4 gamma chain] | C4A CO4 CPAMD2 | Y.IYGK#PVQGVAYVR.F |
| P0C0L4 | Complement C4-A (Acidic complement C4) (C3 and PZP-like alpha-2-macroglobulin domain-containing protein 2) [Cleaved into: Complement C4 beta chain; Complement C4-A alpha chain; C4a anaphylatoxin; C4b-A; C4d-A; Complement C4 gamma chain] | C4A CO4 CPAMD2 | Y.LDK#TEQWSTLPPETK.D |
| P0C0L4 | Complement C4-A (Acidic complement C4) (C3 and PZP-like alpha-2-macroglobulin domain-containing protein 2) [Cleaved into: Complement C4 beta chain; Complement C4-A alpha chain; C4a anaphylatoxin; C4b-A; C4d-A; Complement C4 gamma chain] | C4A CO4 CPAMD2 | Y.LDK#TEQWSTLPPETKDHAVDLIQK.G |
| P0C0L5 | Complement C4-B (Basic complement C4) (C3 and PZP-like alpha-2-macroglobulin domain-containing protein 3) [Cleaved into: Complement C4 beta chain; Complement C4-B alpha chain; C4a anaphylatoxin; C4b-B; C4d-B; Complement C4 gamma chain] | C4B CO4 CPAMD3; C4B_2 | R.GVAHNNLMAMAQETGDNLY*W*GSVTGSQSNAVSPTPAPR#NPSDPM*P.Q |
| P0C0L5 | Complement C4-B (Basic complement C4) (C3 and PZP-like alpha-2-macroglobulin domain-containing protein 3) [Cleaved into: Complement C4 beta chain; Complement C4-B alpha chain; C4a anaphylatoxin; C4b-B; C4d-B; Complement C4 gamma chain] | C4B CO4 CPAMD3; C4B_2 | R.GVAHNNLMAMAQETGDNLYWGSVTGSQSNAVSPTPAPR#NPSDP.M |
| P04003 | C4b-binding protein alpha chain (C4bp) (Proline-rich protein) (PRP) | C4BPA C4BP | C.HPGYK#PTTDEPTTVICQK.N |
| P01031 | Complement C5 (C3 and PZP-like alpha-2-macroglobulin domain-containing protein 4) [Cleaved into: Complement C5 beta chain; Complement C5 alpha chain; C5a anaphylatoxin; Complement C5 alpha' chain] | C5 CPAMD4 | E.DLK#DDQKEMMQTAMQNTMLINGIAQVTFDSETAVK.E |
| P01031 | Complement C5 (C3 and PZP-like alpha-2-macroglobulin domain-containing protein 4) [Cleaved into: Complement C5 beta chain; Complement C5 alpha chain; C5a anaphylatoxin; Complement C5 alpha' chain] | C5 CPAMD4 | K.K#C*C*Y*DGACVNNDETCEQR.A |
| P01031 | Complement C5 (C3 and PZP-like alpha-2-macroglobulin domain-containing protein 4) [Cleaved into: Complement C5 beta chain; Complement C5 alpha chain; C5a anaphylatoxin; Complement C5 alpha' chain] | C5 CPAMD4 | Y.K#EDFSTTGTAYFEVK.E |
| P13671 | Complement component C6 | C6 | K.SNAVDGQWGCWSSWSTCDATYK#.R |
| P13671 | Complement component C6 | C6 | K.SNAVDGQWGCWSSWSTCDATYK#R.S |
| P10643 | Complement component C7 | C7 | A.SSPVNCQW*DFYAPWSECNGCTK#.T |
| P10643 | Complement component C7 | C7 | A.SSPVNCQWDFYAPWSEC*NGCTK#.T |
| P10643 | Complement component C7 | C7 | A.SSPVNCQWDFYAPWSECNGC*TK#.T |
| P10643 | Complement component C7 | C7 | A.SSPVNCQWDFYAPWSECNGCTK#.T |
| P10643 | Complement component C7 | C7 | C.EDSER#RPSCDIDKPPPNIELTGNGYNELTGQFR.N |
| P10643 | Complement component C7 | C7 | C.VCK#MPYECGPSLDVCAQDER.S |
| P10643 | Complement component C7 | C7 | K.ACGAC*PLW*GK#C*DAESSK.C |

| Accession | Protein names | Gene | Peptide |
|---|---|---|---|
| P07357 | Complement component C8 alpha chain (Complement component 8 subunit alpha) | C8A | R.AATPAAVTCQLSNW*SEWTDC*FPC*QDK#K.Y |
| P07357 | Complement component C8 alpha chain (Complement component 8 subunit alpha) | C8A | R.AATPAAVTCQLSNWSEW*TDCFPCQDK#K.Y |
| P07357 | Complement component C8 alpha chain (Complement component 8 subunit alpha) | C8A | R.AATPAAVTCQLSNWSEWTDC*FPCQDK#K.Y |
| P07357 | Complement component C8 alpha chain (Complement component 8 subunit alpha) | C8A | R.AATPAAVTCQLSNWSEWTDCFPC*QDK#K.Y |
| P07357 | Complement component C8 alpha chain (Complement component 8 subunit alpha) | C8A | R.AATPAAVTCQLSNWSEWTDCFPCQDK#.K |
| P07357 | Complement component C8 alpha chain (Complement component 8 subunit alpha) | C8A | R.AATPAAVTCQLSNWSEWTDCFPCQDK#K#.Y |
| P07357 | Complement component C8 alpha chain (Complement component 8 subunit alpha) | C8A | R.AATPAAVTCQLSNWSEWTDCFPCQDK#K.Y |
| P07357 | Complement component C8 alpha chain (Complement component 8 subunit alpha) | C8A | R.AATPAAVTCQLSNWSEWTDCFPCQDK#KY.R |
| P07357 | Complement component C8 alpha chain (Complement component 8 subunit alpha) | C8A | R.AATPAAVTCQLSNWSEWTDCFPCQDKK#Y.R |
| P07357 | Complement component C8 alpha chain (Complement component 8 subunit alpha) | C8A | R.LGSLGAACEQTQTEGAK#ADGSWSCWSSWSVCR#.A |
| P07358 | Complement component C8 beta chain (Complement component 8 subunit beta) | C8B | K.NTPIDGK#WNCWSNWSSCSGR.R |
| P07358 | Complement component C8 beta chain (Complement component 8 subunit beta) | C8B | R.SVDVTLM*PIDCELSSWSSWTTCDPCQK#.K |
| P07358 | Complement component C8 beta chain (Complement component 8 subunit beta) | C8B | R.SVDVTLMPIDCELSSWSSWTTCDPCQK#.K |
| P02748 | Complement component C9 [Cleaved into: Complement component C9a; Complement component C9b] | C9 | C.LCACPFK#FEGIACEISKQK.I |
| P02748 | Complement component C9 [Cleaved into: Complement component C9a; Complement component C9b] | C9 | K.C*LC*AC*PFK#FEGIACEISK.Q |
| P02748 | Complement component C9 [Cleaved into: Complement component C9a; Complement component C9b] | C9 | K.CHTCQNGGTVILMDGK#.C |
| P02748 | Complement component C9 [Cleaved into: Complement component C9a; Complement component C9b] | C9 | R.KCHTCQNGGTVILMDGK#.C |
| Q9NPY3 | Complement component C1q receptor (C1q/MBL/SPA receptor) (C1qR) (C1qR(p)) (C1qRp) (CDw93) (Complement component 1 q subcomponent receptor 1) (Matrix-remodeling-associated protein 4) (CD antigen CD93) | CD93 C1QR1 MXRA4 | R.LLDDLVTCASRNPCSSSPCR#.G |
| P55290 | Cadherin-13 (Heart cadherin) (H-cadherin) (P105) (Truncated cadherin) (T-cad) (T-cadherin) | CDH13 CDHH | R.EGPYIGHVM*EGSPTGTTVM*R#.M |
| P55290 | Cadherin-13 (Heart cadherin) (H-cadherin) (P105) (Truncated cadherin) (T-cad) (T-cadherin) | CDH13 CDHH | R.EGPYIGHVM*EGSPTGTTVMR#.M |
| P55290 | Cadherin-13 (Heart cadherin) (H-cadherin) (P105) (Truncated cadherin) (T-cad) (T-cadherin) | CDH13 CDHH | R.EGPYIGHVMEGSPTGTTVM*R#.M |
| P55290 | Cadherin-13 (Heart cadherin) (H-cadherin) (P105) (Truncated cadherin) (T-cad) (T-cadherin) | CDH13 CDHH | R.EGPYIGHVMEGSPTGTTVMR#.M |
| P55290 | Cadherin-13 (Heart cadherin) (H-cadherin) (P105) (Truncated cadherin) (T-cad) (T-cadherin) | CDH13 CDHH | R.M*TAFDADDPATDNALLR#.Y |
| P55290 | Cadherin-13 (Heart cadherin) (H-cadherin) (P105) (Truncated cadherin) (T-cad) (T-cadherin) | CDH13 CDHH | R.MTAFDADDPATDNALLR#.Y |
| P33151 | Cadherin-5 (7B4 antigen) (Vascular endothelial cadherin) (VE-cadherin) (CD antigen CD144) | CDH5 | K.VCENAVHGQLVLQISAIDK#DITPR.N |
| P33151 | Cadherin-5 (7B4 antigen) (Vascular endothelial cadherin) (VE-cadherin) (CD antigen CD144) | CDH5 | K.VCENAVHGQLVLQISAIDKDITPR#.N |
| P33151 | Cadherin-5 (7B4 antigen) (Vascular endothelial cadherin) (VE-cadherin) (CD antigen CD144) | CDH5 | R.VGTSVGSLFVEDPDEPQNR#.M |
| P00751 | Complement factor B (EC 3.4.21.47) (C3/C5 convertase) (Glycine-rich beta glycoprotein) (GBG) (PBF2) (Properdin factor B) [Cleaved into: Complement factor B Ba fragment; Complement factor B Bb fragment] | CFB BF BFD | K.QLNEINYEDHK#LK.S |
| P00751 | Complement factor B (EC 3.4.21.47) (C3/C5 convertase) (Glycine-rich beta glycoprotein) (GBG) (PBF2) (Properdin factor B) [Cleaved into: Complement factor B Ba fragment; Complement factor B Bb fragment] | CFB BF BFD | K.VSEADSSNADWVTK#QLNEINYEDHK.L |
| P00751 | Complement factor B (EC 3.4.21.47) (C3/C5 convertase) (Glycine-rich beta glycoprotein) (GBG) (PBF2) (Properdin factor B) [Cleaved into: Complement factor B Ba fragment; Complement factor B Bb fragment] | CFB BF BFD | T.TPWSLAR#PQGSCSLEGVEIK.G |

| Accession | Protein names | Gene | Peptide |
|---|---|---|---|
| P00751 | Complement factor B (EC 3.4.21.47) (C3/C5 convertase) (Glycine-rich beta glycoprotein) (GBG) (PBF2) (Properdin factor B) [Cleaved into: Complement factor B Ba fragment; Complement factor B Bb fragment] | CFB BF BFD | T.TTPWSLAR#PQGSCSLEGVEIK.G |
| P08603 | Complement factor H (H factor 1) | CFH HF HF1 HF2 | C.NMGYEY*SER#GDAVCTESGWRPLPSCEEK.S |
| P08603 | Complement factor H (H factor 1) | CFH HF HF1 HF2 | C.NMGYEYSER#GDAVCTESGW*RPLPSCEEK.S |
| P08603 | Complement factor H (H factor 1) | CFH HF HF1 HF2 | C.NMGYEYSER#GDAVCTESGWRPLPSCEEK.S |
| P08603 | Complement factor H (H factor 1) | CFH HF HF1 HF2 | C.NSGY*K#IEGDEEMHCSDDGFWSKEKPK.C |
| P08603 | Complement factor H (H factor 1) | CFH HF HF1 HF2 | K.CGPPPPIDNGDITSFPLSVYAPASSVEY*QC*QNLY*QLEGNK#.R |
| P08603 | Complement factor H (H factor 1) | CFH HF HF1 HF2 | R.DGEKVSVLCQENY*LIQEGEEITC*K#DG.R |
| P08603 | Complement factor H (H factor 1) | CFH HF HF1 HF2 | R.DTSCVNPPTVQNAYIVSRQM*SK#Y*PSGER#VR.Y |
| P08603 | Complement factor H (H factor 1) | CFH HF HF1 HF2 | R.NTEILTGSWSDQTY*PEGTQAIYK#CR#PGYRSLGNVIMVCRKGEWV.A |
| P08603 | Complement factor H (H factor 1) | CFH HF HF1 HF2 | R.NTEILTGSWSDQTYPEGTQAIYK#CRPGYRSLGNVIMVCRKGEWVAL.N |
| P08603 | Complement factor H (H factor 1) | CFH HF HF1 HF2 | R.RNTEILTGSWSDQTYPEGTQAIY*KCR#PGYRSLGNV.I |
| P08603 | Complement factor H (H factor 1) | CFH HF HF1 HF2 | R.TGESVEFVCK#.R |
| P05156 | Complement factor I (EC 3.4.21.45) (C3B/C4B inactivator) [Cleaved into: Complement factor I heavy chain; Complement factor I light chain] | CFI IF | K.ACDGINDCGDQSDELC*C*K#AC*QGK.G |
| P05156 | Complement factor I (EC 3.4.21.45) (C3B/C4B inactivator) [Cleaved into: Complement factor I heavy chain; Complement factor I light chain] | CFI IF | R.CIEGTCVC*K#LPY*QC*PK.N |
| P27918 | Properdin (Complement factor P) | CFP PFC | R.THICNTAVPCPVDGEWDSW*GEW*SPC*IR#RN.M |
| P10909 | Clusterin (Aging-associated gene 4 protein) (Apolipoprotein J) (Apo-J) (Complement cytolysis inhibitor) (CLI) (Complement-associated protein SP-40,40) (Ku70-binding protein 1) (NA1/NA2) (Sulfated glycoprotein 2) (SGP-2) (Testosterone-repressed prostate me | CLU APOJ CLI KUB1 AAG4 | K.LFDSDPITVTVPVEVSRK#NPK#FM*ETVAEK#ALQEYRKKH.R |
| P10909 | Clusterin (Aging-associated gene 4 protein) (Apolipoprotein J) (Apo-J) (Complement cytolysis inhibitor) (CLI) (Complement-associated protein SP-40,40) (Ku70-binding protein 1) (NA1/NA2) (Sulfated glycoprotein 2) (SGP-2) (Testosterone-repressed prostate me | CLU APOJ CLI KUB1 AAG4 | R.EILSVDCSTNNPSQAK#L.R |
| P10909 | Clusterin (Aging-associated gene 4 protein) (Apolipoprotein J) (Apo-J) (Complement cytolysis inhibitor) (CLI) (Complement-associated protein SP-40,40) (Ku70-binding protein 1) (NA1/NA2) (Sulfated glycoprotein 2) (SGP-2) (Testosterone-repressed prostate me | CLU APOJ CLI KUB1 AAG4 | R.M*KDQC*DKC*R#EILSVDCSTNNPSQAK.L |
| P10909 | Clusterin (Aging-associated gene 4 protein) (Apolipoprotein J) (Apo-J) (Complement cytolysis inhibitor) (CLI) (Complement-associated protein SP-40,40) (Ku70-binding protein 1) (NA1/NA2) (Sulfated glycoprotein 2) (SGP-2) (Testosterone-repressed prostate me | CLU APOJ CLI KUB1 AAG4 | Y.VNK#EIQNAVNGVK.Q |
| P39060 | Collagen alpha-1(XVIII) chain [Cleaved into: Endostatin; Non-collagenous domain 1 (NC1)] | COL18A1 | R.AAVPIVNLKDELLFPSW*EALFSGSEGPLK#PGA.R |
| P00450 | Ceruloplasmin (EC 1.16.3.1) (Ferroxidase) | CP | K.ERGPEEEHLGILGPVIW*AEVGDTIR#VTFHNK#GAY*PLSIEPIG.V |
| P00450 | Ceruloplasmin (EC 1.16.3.1) (Ferroxidase) | CP | K.ERGPEEEHLGILGPVIWAEVGDTIR#VTFHNK#GAY*PLSIEPIGVRFN.K |
| P00450 | Ceruloplasmin (EC 1.16.3.1) (Ferroxidase) | CP | K.HIDREFVVMFSVVDENFSWYLEDNIK#TY*C*SEP.E |
| P00450 | Ceruloplasmin (EC 1.16.3.1) (Ferroxidase) | CP | K.HYYIGIIETTWDY*ASDHGEKK#LISVDTEHS.N |
| P00450 | Ceruloplasmin (EC 1.16.3.1) (Ferroxidase) | CP | K.NNEGTYYSPNY*NPQSR#SV.P |
| P00450 | Ceruloplasmin (EC 1.16.3.1) (Ferroxidase) | CP | R.MFTTAPDQVDKEDEDFQESNK#M*HSM*NGFMY*GNQPGLT.M |
| P00450 | Ceruloplasmin (EC 1.16.3.1) (Ferroxidase) | CP | R.MFTTAPDQVDKEDEDFQESNK#MHSMNGFM*Y*GNQPGLT.M |
| P00450 | Ceruloplasmin (EC 1.16.3.1) (Ferroxidase) | CP | R.SGAGTEDSACIPW*AY*Y*STVDQVK#.D |

| Accession | Protein names | Gene | Peptide |
|---|---|---|---|
| P00450 | Ceruloplasmin (EC 1.16.3.1) (Ferroxidase) | CP | R.SGAGTEDSACIPWAYYSTVDQVK#DLYSGLIGP.L |
| P00450 | Ceruloplasmin (EC 1.16.3.1) (Ferroxidase) | CP | Y.CSEPEK#VDK#DNEDFQESNR.M |
| Q9NQ79 | Cartilage acidic protein 1 (68 kDa chondrocyte-expressed protein) (CEP-68) (ASPIC) | CRTAC1 ASPIC1 CEP68 | R.DKPVCVNTY*GSYR#.C |
| Q08554 | Desmocollin-1 (Cadherin family member 1) (Desmosomal glycoprotein 2/3) (DG2/DG3) | DSC1 CDHF1 | R.SGTSVGKVTATDLDEPDTLHTR#.L |
| Q02487 | Desmocollin-2 (Cadherin family member 2) (Desmocollin-3) (Desmosomal glycoprotein II) (Desmosomal glycoprotein III) | DSC2 CDHF2 DSC3 | R.VGTTVGQVCATDK#DEPDTM*HTR.L |
| Q13822 | Ectonucleotide pyrophosphatase/phosphodiesterase family member 2 (E-NPP 2) (EC 3.1.4.39) (Autotaxin) (Extracellular lysophospholipase D) (LysoPLD) | ENPP2 ATX PDNP2 | C.VNVIFVGDHGMEDVTC*DR#TEFLSNYLTNVDDITLVPGTLGR.I |
| Q13822 | Ectonucleotide pyrophosphatase/phosphodiesterase family member 2 (E-NPP 2) (EC 3.1.4.39) (Autotaxin) (Extracellular lysophospholipase D) (LysoPLD) | ENPP2 ATX PDNP2 | C.VNVIFVGDHGMEDVTCDR#TEFLSNYLTNVDDITLVPGTLGR.I |
| P00742 | Coagulation factor X (EC 3.4.21.6) (Stuart factor) (Stuart-Prower factor) [Cleaved into: Factor X light chain; Factor X heavy chain; Activated factor Xa heavy chain] | F10 | K.LCSLDNGDCDQFCHEEQNSVVCSCAR#.G |
| P00742 | Coagulation factor X (EC 3.4.21.6) (Stuart factor) (Stuart-Prower factor) [Cleaved into: Factor X light chain; Factor X heavy chain; Activated factor Xa heavy chain] | F10 | R.KLCSLDNGDCDQFC*HEEQNSVVCSCAR#.G |
| P00742 | Coagulation factor X (EC 3.4.21.6) (Stuart factor) (Stuart-Prower factor) [Cleaved into: Factor X light chain; Factor X heavy chain; Activated factor Xa heavy chain] | F10 | R.KLCSLDNGDCDQFCHEEQNSVVCSCAR#.G |
| P00748 | Coagulation factor XII (EC 3.4.21.38) (Hageman factor) (HAF) [Cleaved into: Coagulation factor XIIa heavy chain; Beta-factor XIIa part 1; Coagulation factor XIIa light chain (Beta-factor XIIa part 2)] | F12 | R.LQEDADGSCALLSPYVQPVCLPSGAAR#PSETTL.C |
| P05160 | Coagulation factor XIII B chain (Fibrin-stabilizing factor B subunit) (Protein-glutamine gamma-glutamyltransferase B chain) (Transglutaminase B chain) | F13B | C.EQGK#WSSPPVCLEPCTVNVDYMNR.N |
| P05160 | Coagulation factor XIII B chain (Fibrin-stabilizing factor B subunit) (Protein-glutamine gamma-glutamyltransferase B chain) (Transglutaminase B chain) | F13B | C.LAGYTTESGR#QEEQTTCTTEGWSPEPR.C |
| P00734 | Prothrombin (EC 3.4.21.5) (Coagulation factor II) [Cleaved into: Activation peptide fragment 1; Activation peptide fragment 2; Thrombin light chain; Thrombin heavy chain] | F2 | K.HQDFNSAVQLVENFCR#NPDGDEEGVW.C |
| P00734 | Prothrombin (EC 3.4.21.5) (Coagulation factor II) [Cleaved into: Activation peptide fragment 1; Activation peptide fragment 2; Thrombin light chain; Thrombin heavy chain] | F2 | R.IVEGSDAEIGMSPWQVMLFR#K#SPQELLCGASLISDRW.V |
| P00734 | Prothrombin (EC 3.4.21.5) (Coagulation factor II) [Cleaved into: Activation peptide fragment 1; Activation peptide fragment 2; Thrombin light chain; Thrombin heavy chain] | F2 | R.IVEGSDAEIGMSPWQVMLFRK#SPQ.E |
| P00734 | Prothrombin (EC 3.4.21.5) (Coagulation factor II) [Cleaved into: Activation peptide fragment 1; Activation peptide fragment 2; Thrombin light chain; Thrombin heavy chain] | F2 | Y.NW*R#ENLDRDIALMK.L |
| P12259 | Coagulation factor V (Activated protein C cofactor) (Proaccelerin, labile factor) [Cleaved into: Coagulation factor V heavy chain; Coagulation factor V light chain] | F5 | R.DIASGLIGLLLICK#SR.S |
| P23142 | Fibulin-1 (FIBL-1) | FBLN1 PP213 | H.TCINTEGSYTCQK#.N |
| P23142 | Fibulin-1 (FIBL-1) | FBLN1 PP213 | K.C*ENTLGSYLCSCSVGFR#.L |
| P23142 | Fibulin-1 (FIBL-1) | FBLN1 PP213 | K.CENTLGSYLCSCSVGFR#.L |
| P23142 | Fibulin-1 (FIBL-1) | FBLN1 PP213 | K.DIDECESGIHNCLPDFIC*QNTLGSFR#.C |
| P23142 | Fibulin-1 (FIBL-1) | FBLN1 PP213 | R.C*INIPGSFQCSCPSSGYR#.L |
| P23142 | Fibulin-1 (FIBL-1) | FBLN1 PP213 | R.CINIPGSFQCSCPSSGYR#.L |
| P23142 | Fibulin-1 (FIBL-1) | FBLN1 PP213 | R.DSSCGTGYELTEDNSCK#DIDECESGIHNCLPDFIC*QNTLG.S |
| P23142 | Fibulin-1 (FIBL-1) | FBLN1 PP213 | R.LCGHKC*ENTLGSYLCSCSVGFR#.L |
| P23142 | Fibulin-1 (FIBL-1) | FBLN1 PP213 | R.LGESC*INTVGSFR#.C |
| P23142 | Fibulin-1 (FIBL-1) | FBLN1 PP213 | R.LGESCINTVGSFR#.C |
| P98095 | Fibulin-2 (FIBL-2) | FBLN2 | K.GSFYCQAR#.Q |
| P35555 | Fibrillin-1 [Cleaved into: Asprosin] | FBN1 FBN | R.CVNTDGSYR#.C |
| P35555 | Fibrillin-1 [Cleaved into: Asprosin] | FBN1 FBN | R.KCAPGTCQNLDGSYR#.C |
| P35555 | Fibrillin-1 [Cleaved into: Asprosin] | FBN1 FBN | R.NAECINTAGSYR#.C |
| P35555 | Fibrillin-1 [Cleaved into: Asprosin] | FBN1 FBN | R.NTIGSFNCR#.C |

| Accession | Protein names | Gene | Peptide |
|---|---|---|---|
| P02675 | Fibrinogen beta chain [Cleaved into: Fibrinopeptide B; Fibrinogen beta chain] | FGB | R.KAPDAGGCLHADPDLGVLC*PTGC*QLQEALLQQER#PI.R |
| P02751 | Fibronectin (FN) (Cold-insoluble globulin) (CIG) [Cleaved into: Anastellin; Ugl-Y1; Ugl-Y2; Ugl-Y3] | FN1 FN | C.DNCR#RPGGEPSPEGTTGQSYNQYSQR.Y |
| P02751 | Fibronectin (FN) (Cold-insoluble globulin) (CIG) [Cleaved into: Anastellin; Ugl-Y1; Ugl-Y2; Ugl-Y3] | FN1 FN | K.CFDHAAGTSYVVGETWEK#PY*QGW*M*MVDCTCLGEGSGR.I |
| P02751 | Fibronectin (FN) (Cold-insoluble globulin) (CIG) [Cleaved into: Anastellin; Ugl-Y1; Ugl-Y2; Ugl-Y3] | FN1 FN | R.GFNCESKPEAEETC*FDK#Y*TGNTY*R.V |
| P02751 | Fibronectin (FN) (Cold-insoluble globulin) (CIG) [Cleaved into: Anastellin; Ugl-Y1; Ugl-Y2; Ugl-Y3] | FN1 FN | R.QGENGQM*M*SCTC*LGNGK#G.E |
| P02751 | Fibronectin (FN) (Cold-insoluble globulin) (CIG) [Cleaved into: Anastellin; Ugl-Y1; Ugl-Y2; Ugl-Y3] | FN1 FN | R.QGENGQMM*SCTC*LGNGK#G.E |
| P02751 | Fibronectin (FN) (Cold-insoluble globulin) (CIG) [Cleaved into: Anastellin; Ugl-Y1; Ugl-Y2; Ugl-Y3] | FN1 FN | R.QGENGQMMSCTC*LGNGK#G.E |
| P02751 | Fibronectin (FN) (Cold-insoluble globulin) (CIG) [Cleaved into: Anastellin; Ugl-Y1; Ugl-Y2; Ugl-Y3] | FN1 FN | R.RPGGEPSPEGTTGQSYNQYSQR#.Y |
| P02751 | Fibronectin (FN) (Cold-insoluble globulin) (CIG) [Cleaved into: Anastellin; Ugl-Y1; Ugl-Y2; Ugl-Y3] | FN1 FN | R.TYLGNALVC*TC*Y*GGSR#.G |
| P02751 | Fibronectin (FN) (Cold-insoluble globulin) (CIG) [Cleaved into: Anastellin; Ugl-Y1; Ugl-Y2; Ugl-Y3] | FN1 FN | R.W*K#C*DPVDQC*QDSETGTFYQIGDSWEK.Y |
| P02774 | Vitamin D-binding protein (DBP) (VDB) (Gc protein-derived macrophage activating factor) (Gc-MAF) (GcMAF) (Gc-globulin) (Group-specific component) (Gc) (Vitamin D-binding protein-macrophage activating factor) (DBP-maf) | GC | C.CESASEDCM*AK#ELPEHTVKLCDNLSTK.N |
| P02774 | Vitamin D-binding protein (DBP) (VDB) (Gc protein-derived macrophage activating factor) (Gc-MAF) (GcMAF) (Gc-globulin) (Group-specific component) (Gc) (Vitamin D-binding protein-macrophage activating factor) (DBP-maf) | GC | C.CESASEDCMAK#ELPEHTVKLCDNLSTK.N |
| P02774 | Vitamin D-binding protein (DBP) (VDB) (Gc protein-derived macrophage activating factor) (Gc-MAF) (GcMAF) (Gc-globulin) (Group-specific component) (Gc) (Vitamin D-binding protein-macrophage activating factor) (DBP-maf) | GC | K.EFSHLGKEDFTSLSLVLYSR#K#FPSGTFEQVSQLVKEVV.S |
| P02774 | Vitamin D-binding protein (DBP) (VDB) (Gc protein-derived macrophage activating factor) (Gc-MAF) (GcMAF) (Gc-globulin) (Group-specific component) (Gc) (Vitamin D-binding protein-macrophage activating factor) (DBP-maf) | GC | K.ELSSFIDKGQELC*ADY*SENTFTEY*K#.K |
| P02774 | Vitamin D-binding protein (DBP) (VDB) (Gc protein-derived macrophage activating factor) (Gc-MAF) (GcMAF) (Gc-globulin) (Group-specific component) (Gc) (Vitamin D-binding protein-macrophage activating factor) (DBP-maf) | GC | K.EVVSLTEACC*AEGADPDC*Y*DTR#.T |
| P02774 | Vitamin D-binding protein (DBP) (VDB) (Gc protein-derived macrophage activating factor) (Gc-MAF) (GcMAF) (Gc-globulin) (Group-specific component) (Gc) (Vitamin D-binding protein-macrophage activating factor) (DBP-maf) | GC | K.EVVSLTEACCAEGADPDC*Y*DTR#TSALSAK.S |
| P02774 | Vitamin D-binding protein (DBP) (VDB) (Gc protein-derived macrophage activating factor) (Gc-MAF) (GcMAF) (Gc-globulin) (Group-specific component) (Gc) (Vitamin D-binding protein-macrophage activating factor) (DBP-maf) | GC | K.EVVSLTEACCAEGADPDCY*DTR#TSALSAK#.S |
| P02774 | Vitamin D-binding protein (DBP) (VDB) (Gc protein-derived macrophage activating factor) (Gc-MAF) (GcMAF) (Gc-globulin) (Group-specific component) (Gc) (Vitamin D-binding protein-macrophage activating factor) (DBP-maf) | GC | K.HQPQEFPTYVEPTNDEIC*EAFRK#DPKEY*ANQ.F |
| P02774 | Vitamin D-binding protein (DBP) (VDB) (Gc protein-derived macrophage activating factor) (Gc-MAF) (GcMAF) (Gc-globulin) (Group-specific component) (Gc) (Vitamin D-binding protein-macrophage activating factor) (DBP-maf) | GC | K.HQPQEFPTYVEPTNDEICEAFRK#DPK#EYAN.Q |
| P02774 | Vitamin D-binding protein (DBP) (VDB) (Gc protein-derived macrophage activating factor) (Gc-MAF) (GcMAF) (Gc-globulin) (Group-specific component) (Gc) (Vitamin D-binding protein-macrophage activating factor) (DBP-maf) | GC | K.LAQKVPTADLEDVLPLAEDITNILSK#.C |
| P02774 | Vitamin D-binding protein (DBP) (VDB) (Gc protein-derived macrophage activating factor) (Gc-MAF) (GcMAF) (Gc-globulin) (Group-specific component) (Gc) (Vitamin D-binding protein-macrophage activating factor) (DBP-maf) | GC | K.LAQKVPTADLEDVLPLAEDITNILSK#C*C*ESASEDC*MAK.E |
| P02774 | Vitamin D-binding protein (DBP) (VDB) (Gc protein-derived macrophage activating factor) (Gc-MAF) (GcMAF) (Gc-globulin) (Group-specific component) (Gc) (Vitamin D-binding protein-macrophage activating factor) (DBP-maf) | GC | K.SCESNSPFPVHPGTAECCTKEGLERK#LC*MAALKHQPQEF.P |
| P06396 | Gelsolin (AGEL) (Actin-depolymerizing factor) (ADF) (Brevin) | GSN | N.WR#DPDQTDGLGLSYLSSHIANVER.V |
| P06396 | Gelsolin (AGEL) (Actin-depolymerizing factor) (ADF) (Brevin) | GSN | R.FVIEEVPGELM*QEDLATDDVMLLDTWDQVFVWVGK#DSQEEEKTEAL.T |
| P06396 | Gelsolin (AGEL) (Actin-depolymerizing factor) (ADF) (Brevin) | GSN | R.FVIEEVPGELMQEDLATDDVM*LLDTWDQVFVWVGK#DSQEEEKTEAL.T |

| Accession | Protein names | Gene | Peptide |
|---|---|---|---|
| P06396 | Gelsolin (AGEL) (Actin-depolymerizing factor) (ADF) (Brevin) | GSN | R.FVIEEVPGELMQEDLATDDVMLLDTW*DQVFVWVGK#DSQEEEKTEAL.T |
| P09211 | Glutathione S-transferase P (EC 2.5.1.18) (GST class-pi) (GSTP1-1) | GSTP1 FAEES3 GST3 | R.MLLADQGQSWKEEVVTVETW*QEGSLK#ASC*LY*GQLPKFQ.D |
| P68871 | Hemoglobin subunit beta (Beta-globin) (Hemoglobin beta chain) [Cleaved into: LVV-hemorphin-7; Spinorphin] | HBB | A.VTALWGK#VNVDEVGGEALGR.L |
| P02042 | Hemoglobin subunit delta (Delta-globin) (Hemoglobin delta chain) | HBD | K.K#VLGAFSDGLAHLDNLK.G |
| P02008 | Hemoglobin subunit zeta (HBAZ) (Hemoglobin zeta chain) (Zeta-globin) | HBZ HBZ2 | K.LLSHCLLVTLAAR#.F |
| P02790 | Hemopexin (Beta-1B-glycoprotein) | HPX | K.EVGTPHGIILDSVDAAFIC*PGSSR#LH.I |
| P02790 | Hemopexin (Beta-1B-glycoprotein) | HPX | K.EVGTPHGIILDSVDAAFICPGSSR#LHIMAGR#R#LW.W |
| P02790 | Hemopexin (Beta-1B-glycoprotein) | HPX | K.EVGTPHGIILDSVDAAFICPGSSRLHIM*AGR#R#LWWLDLK#SGAQATWT.E |
| P02790 | Hemopexin (Beta-1B-glycoprotein) | HPX | K.EVGTPHGIILDSVDAAFICPGSSRLHIMAGR#R#LWWLDLK#SG.A |
| P02790 | Hemopexin (Beta-1B-glycoprotein) | HPX | K.SGAQATWTELPWPHEK#VDGALC*.M |
| P02790 | Hemopexin (Beta-1B-glycoprotein) | HPX | K.SGAQATWTELPWPHEK#VDGALC.M |
| P02790 | Hemopexin (Beta-1B-glycoprotein) | HPX | K.SLGPNSCSANGPGLYLIHGPNLYCY*SDVEK#L.N |
| P02790 | Hemopexin (Beta-1B-glycoprotein) | HPX | R.GECQAEGVLFFQGDR#EWFW*DLATGTM*K#.E |
| P02790 | Hemopexin (Beta-1B-glycoprotein) | HPX | R.GECQAEGVLFFQGDREWFWDLATGTMK#ERSW*PA.V |
| P02790 | Hemopexin (Beta-1B-glycoprotein) | HPX | R.WK#NFPSPVDAAFR.Q |
| P04196 | Histidine-rich glycoprotein (Histidine-proline-rich glycoprotein) (HPRG) | HRG | R.KYWNDCEPPDSR#R.P |
| P04196 | Histidine-rich glycoprotein (Histidine-proline-rich glycoprotein) (HPRG) | HRG | Y.K#EENDDFASFRVDR.I |
| P01344 | Insulin-like growth factor II (IGF-II) (Somatomedin-A) (T3M-11-derived growth factor) [Cleaved into: Insulin-like growth factor II; Insulin-like growth factor II Ala-25 Del; Preptin] | IGF2 PP1446 | Y.R#PSETLC*GGELVDTLQFVCGDR.G |
| P05106 | Integrin beta-3 (Platelet membrane glycoprotein IIIa) (GPIIIa) (CD antigen CD61) | ITGB3 GP3A | C.DLK#ENLLKDNCAPESIEFPVSEAR.V |
| P19827 | Inter-alpha-trypsin inhibitor heavy chain H1 (ITI heavy chain H1) (ITI-HC1) (Inter-alpha-inhibitor heavy chain 1) (Inter-alpha-trypsin inhibitor complex component III) (Serum-derived hyaluronan-associated protein) (SHAP) | ITIH1 IGHEP1 | K.QLVHHFEIDVDIFEPQGISK#L.D |
| P19827 | Inter-alpha-trypsin inhibitor heavy chain H1 (ITI heavy chain H1) (ITI-HC1) (Inter-alpha-inhibitor heavy chain 1) (Inter-alpha-trypsin inhibitor complex component III) (Serum-derived hyaluronan-associated protein) (SHAP) | ITIH1 IGHEP1 | R.IYEDHDATQQLQGFYSQVAK#PLLVDVDLQY*PQDAVLALTQN.H |
| P19823 | Inter-alpha-trypsin inhibitor heavy chain H2 (ITI heavy chain H2) (ITI-HC2) (Inter-alpha-inhibitor heavy chain 2) (Inter-alpha-trypsin inhibitor complex component II) (Serum-derived hyaluronan-associated protein) (SHAP) | ITIH2 IGHEP2 | K.AGELEVFNGYFVHFFAPDNLDPIPK#NILFVIDVSGSMWG.V |
| P19823 | Inter-alpha-trypsin inhibitor heavy chain H2 (ITI heavy chain H2) (ITI-HC2) (Inter-alpha-inhibitor heavy chain 2) (Inter-alpha-trypsin inhibitor complex component II) (Serum-derived hyaluronan-associated protein) (SHAP) | ITIH2 IGHEP2 | K.FDPAKLDQIESVITATSANTQLVLETLAQMDDLQDFLSK#DKHAD.P |
| P19823 | Inter-alpha-trypsin inhibitor heavy chain H2 (ITI heavy chain H2) (ITI-HC2) (Inter-alpha-inhibitor heavy chain 2) (Inter-alpha-trypsin inhibitor complex component II) (Serum-derived hyaluronan-associated protein) (SHAP) | ITIH2 IGHEP2 | R.LAK#HLEVDVWVIEPQGLR.F |
| P19823 | Inter-alpha-trypsin inhibitor heavy chain H2 (ITI heavy chain H2) (ITI-HC2) (Inter-alpha-inhibitor heavy chain 2) (Inter-alpha-trypsin inhibitor complex component II) (Serum-derived hyaluronan-associated protein) (SHAP) | ITIH2 IGHEP2 | Y.IEK#IQPSGGTNINEALLR.A |
| Q06033 | Inter-alpha-trypsin inhibitor heavy chain H3 (ITI heavy chain H3) (ITI-HC3) (Inter-alpha-inhibitor heavy chain 3) (Serum-derived hyaluronan-associated protein) (SHAP) | ITIH3 | R.LVDEDMNSFKADVK#.G |
| Q06033 | Inter-alpha-trypsin inhibitor heavy chain H3 (ITI heavy chain H3) (ITI-HC3) (Inter-alpha-inhibitor heavy chain 3) (Serum-derived hyaluronan-associated protein) (SHAP) | ITIH3 | R.LWAYLTIEQLLEK#RK#NAHGE.E |
| Q14624 | Inter-alpha-trypsin inhibitor heavy chain H4 (ITI heavy chain H4) (ITI-HC4) (Inter-alpha-inhibitor heavy chain 4) (Inter-alpha-trypsin inhibitor family heavy chain-related protein) (IHRP) (Plasma kallikrein sensitive glycoprotein 120) (Gp120) (PK-120) [Cl | ITIH4 IHRP ITIHL1 PK120 PRO1851 | K.WK#ETLFSVMPGLK.M |
| Q14624 | Inter-alpha-trypsin inhibitor heavy chain H4 (ITI heavy chain H4) (ITI-HC4) (Inter-alpha-inhibitor heavy chain 4) (Inter-alpha-trypsin inhibitor family heavy chain- | ITIH4 IHRP ITIHL1 PK120 PRO1851 | R.LWAYLTIQQLLEQTVSASDADQQALR#.N |

| Accession | Protein names | Gene | Peptide |
|---|---|---|---|
| | related protein) (IHRP) (Plasma kallikrein sensitive glycoprotein 120) (Gp120) (PK-120) [Cl | | |
| P03952 | Plasma kallikrein (EC 3.4.21.34) (Fletcher factor) (Kininogenin) (Plasma prekallikrein) (PKK) [Cleaved into: Plasma kallikrein heavy chain; Plasma kallikrein light chain] | KLKB1 KLK3 | R.GGDVASMYTPNAQY*C*QM*R#.C |
| P03952 | Plasma kallikrein (EC 3.4.21.34) (Fletcher factor) (Kininogenin) (Plasma prekallikrein) (PKK) [Cleaved into: Plasma kallikrein heavy chain; Plasma kallikrein light chain] | KLKB1 KLK3 | R.LCNTGDNSVC*TTK#TS.T |
| P01042 | Kininogen-1 (Alpha-2-thiol proteinase inhibitor) (Fitzgerald factor) (High molecular weight kininogen) (HMWK) (Williams-Fitzgerald-Flaujeac factor) [Cleaved into: Kininogen-1 heavy chain; T-kinin (Ile-Ser-Bradykinin); Bradykinin (Kallidin I); Lysyl-bradyk | KNG1 BDK KNG | K.SLWNGDTGECTDNAYIDIQLR#IASFSQN.C |
| P01042 | Kininogen-1 (Alpha-2-thiol proteinase inhibitor) (Fitzgerald factor) (High molecular weight kininogen) (HMWK) (Williams-Fitzgerald-Flaujeac factor) [Cleaved into: Kininogen-1 heavy chain; T-kinin (Ile-Ser-Bradykinin); Bradykinin (Kallidin I); Lysyl-bradyk | KNG1 BDK KNG | K.SLWNGDTGECTDNAYIDIQLRIASFSQNCDIY*PGK#DFVQPP.T |
| P01042 | Kininogen-1 (Alpha-2-thiol proteinase inhibitor) (Fitzgerald factor) (High molecular weight kininogen) (HMWK) (Williams-Fitzgerald-Flaujeac factor) [Cleaved into: Kininogen-1 heavy chain; T-kinin (Ile-Ser-Bradykinin); Bradykinin (Kallidin I); Lysyl-bradyk | KNG1 BDK KNG | K.TWQDCEYKDAAKAATGECTATVGK#RSST.K |
| P08519 | Apolipoprotein(a) (Apo(a)) (Lp(a)) (EC 3.4.21.-) | LPA | R.TPENYPNAGLTR#NYCRNPDAEIRPW*CYTMDPSVR.W |
| P02750 | Leucine-rich alpha-2-glycoprotein (LRG) | LRG1 LRG | R.DGFDISGNPWICDQNLSDLY*RW*LQAQKDK#MFSQN.D |
| Q07954 | Prolow-density lipoprotein receptor-related protein 1 (LRP-1) (Alpha-2-macroglobulin receptor) (A2MR) (Apolipoprotein E receptor) (APOER) (CD antigen CD91) [Cleaved into: Low-density lipoprotein receptor-related protein 1 85 kDa subunit (LRP-85); Low-dens | LRP1 A2MR APR | K.CDQNKFSVK#.C |
| Q07954 | Prolow-density lipoprotein receptor-related protein 1 (LRP-1) (Alpha-2-macroglobulin receptor) (A2MR) (Apolipoprotein E receptor) (APOER) (CD antigen CD91) [Cleaved into: Low-density lipoprotein receptor-related protein 1 85 kDa subunit (LRP-85); Low-dens | LRP1 A2MR APR | R.VNNGGCSSLCLATPGSR#.Q |
| P02788 | Lactotransferrin (Lactoferrin) (EC 3.4.21.-) (Growth-inhibiting protein 12) (Talalactoferrin) [Cleaved into: Lactoferricin-H (Lfcin-H); Kaliocin-1; Lactoferroxin-A; Lactoferroxin-B; Lactoferroxin-C] | LTF GIG12 LF | C.LAENAGDVAFVK#DVTVLQNTDGNNNEAWAK.D |
| Q04721 | Neurogenic locus notch homolog protein 2 (Notch 2) (hN2) [Cleaved into: Notch 2 extracellular truncation (N2ECD); Notch 2 intracellular domain (N2ICD)] | NOTCH2 | K.AGLLCHLDDACISNPCHK#.G |
| O14786 | Neuropilin-1 (Vascular endothelial cell growth factor 165 receptor) (CD antigen CD304) | NRP1 NRP VEGF165R | Y.DR#LEIW*DGFPDVGPHIGR.Y |
| Q9UN70 | Protocadherin gamma-C3 (PCDH-gamma-C3) (Protocadherin-2) (Protocadherin-43) (PC-43) | PCDHGC3 PCDH2 | R.DAGTPSLSALTIVR#.V |
| Q96PD5 | N-acetylmuramoyl-L-alanine amidase (EC 3.5.1.28) (Peptidoglycan recognition protein 2) (Peptidoglycan recognition protein long) (PGRP-L) | PGLYRP2 PGLYRPL PGRPL UNQ3103/PRO10102 | R.TDCPGDALFDLLR#TWPHFTATVK#.P |
| P14618 | Pyruvate kinase PKM (EC 2.7.1.40) (Cytosolic thyroid hormone-binding protein) (CTHBP) (Opa-interacting protein 3) (OIP-3) (Pyruvate kinase 2/3) (Pyruvate kinase muscle isozyme) (Threonine-protein kinase PKM2) (EC 2.7.11.1) (Thyroid hormone-binding protein | PKM OIP3 PK2 PK3 PKM2 | K.ITLDNAYM*EK#C*DENILWL.D |
| P00747 | Plasminogen (EC 3.4.21.7) [Cleaved into: Plasmin heavy chain A; Activation peptide; Angiostatin; Plasmin heavy chain A, short form; Plasmin light chain B] | PLG | C.LPSPNYVVADR#TECFITGWGETQGTFGAGLLK.E |
| P00747 | Plasminogen (EC 3.4.21.7) [Cleaved into: Plasmin heavy chain A; Activation peptide; Angiostatin; Plasmin heavy chain A, short form; Plasmin light chain B] | PLG | R.FSPATHPSEGLEENYC*R#NPDNDPQGPW*CYTTDPEK.R |
| P00747 | Plasminogen (EC 3.4.21.7) [Cleaved into: Plasmin heavy chain A; Activation peptide; Angiostatin; Plasmin heavy chain A, short form; Plasmin light chain B] | PLG | R.NPDADKGPWCFTTDPSVR#W*EY*C*NLKKC.S |
| P00747 | Plasminogen (EC 3.4.21.7) [Cleaved into: Plasmin heavy chain A; Activation peptide; Angiostatin; Plasmin heavy chain A, short form; Plasmin light chain B] | PLG | R.VQSTELCAGHLAGGTDSCQGDSGGPLVC*FEK#.D |
| P02775 | Platelet basic protein (PBP) (C-X-C motif chemokine 7) (Leukocyte-derived growth factor) (LDGF) (Macrophage-derived growth factor) (MDGF) (Small-inducible cytokine B7) [Cleaved into: Connective tissue-activating peptide III (CTAP-III) (LA-PF4) (Low-affinity | PPBP CTAP3 CXCL7 SCYB7 TGB1 THBGB1 | K.GKEESLDSDLYAELR#C*M*C*IK.T |
| P02753 | Retinol-binding protein 4 (Plasma retinol-binding protein) (PRBP) (RBP) [Cleaved into: Plasma retinol-binding protein(1-182); Plasma retinol-binding protein(1-181); Plasma retinol-binding protein(1-179); Plasma retinol-binding protein(1-176)] | RBP4 PRO2222 | K.KDPEGLFLQDNIVAEFSVDETGQM*SATAK#GR.V |

| Accession | Protein names | Gene | Peptide |
|---|---|---|---|
| P07998 | Ribonuclease pancreatic (EC 4.6.1.18) (HP-RNase) (RIB-1) (RNase UpI-1) (Ribonuclease 1) (RNase 1) (Ribonuclease A) (RNase A) | RNASE1 RIB1 RNS1 | C.K#PVNTFVHEPLVDVQNVCFQEK.V |
| P13521 | Secretogranin-2 (Chromogranin-C) (Secretogranin II) (SgII) [Cleaved into: Secretoneurin (SN); Manserin] | SCG2 CHGC | R.SGQLGIQEEDLR#K#ES.K |
| P01011 | Alpha-1-antichymotrypsin (ACT) (Cell growth-inhibiting gene 24/25 protein) (Serpin A3) [Cleaved into: Alpha-1-antichymotrypsin His-Pro-less] | SERPINA3 AACT GIG24 GIG25 | K.AVLDVFEEGTEASAATAVK#ITLLSALVETR#TIVR#FNR#.P |
| P01011 | Alpha-1-antichymotrypsin (ACT) (Cell growth-inhibiting gene 24/25 protein) (Serpin A3) [Cleaved into: Alpha-1-antichymotrypsin His-Pro-less] | SERPINA3 AACT GIG24 GIG25 | K.RLYGSEAFATDFQDSAAAK#K#LINDYVK#NGTR#.G |
| P01011 | Alpha-1-antichymotrypsin (ACT) (Cell growth-inhibiting gene 24/25 protein) (Serpin A3) [Cleaved into: Alpha-1-antichymotrypsin His-Pro-less] | SERPINA3 AACT GIG24 GIG25 | R.GKITDLIKDLDSQTM*MVLVNYIFFKAK#WEMPFDPQDTH.Q |
| P01008 | Antithrombin-III (ATIII) (Serpin C1) | SERPINC1 AT3 PRO0309 | K.AFLEVNEEGSEAAASTAVVIAGRSLNPNR#.V |
| P01008 | Antithrombin-III (ATIII) (Serpin C1) | SERPINC1 AT3 PRO0309 | K.ELFYKADGESCSASM*M*Y*QEGK#.F |
| P01008 | Antithrombin-III (ATIII) (Serpin C1) | SERPINC1 AT3 PRO0309 | R.FRIEDGFSLKEQLQDMGLVDLFSPEK#SK#.L |
| P08697 | Alpha-2-antiplasmin (Alpha-2-AP) (Alpha-2-plasmin inhibitor) (Alpha-2-PI) (Serpin F2) | SERPINF2 AAP PLI | R.GISEQSLVVSGVQHQSTLELSEVGVEAAAATSIAM*SR#MSLSSFSV.N |
| Q8TER0 | Sushi, nidogen and EGF-like domain-containing protein 1 (Insulin-responsive sequence DNA-binding protein 1) (IRE-BP1) | SNED1 | K.CDCPPGFSGR#.H |
| P07996 | Thrombospondin-1 (Glycoprotein G) | THBS1 TSP TSP1 | K.QDCPIDGCLSNPCFAGVK#.C |
| Q9Y490 | Talin-1 | TLN1 KIAA1027 TLN | R.DDILNGSHPVSFDK#ACEFAGFQ.C |
| Q9H853 | Putative tubulin-like protein alpha-4B (EC 3.6.5.-) (Alpha-tubulin 4B) | TUBA4B TUBA4 | R.QIFHPEQLITGK#EDAANN.Y |
| P54578 | Ubiquitin carboxyl-terminal hydrolase 14 (EC 3.4.19.12) (Deubiquitinating enzyme 14) (Ubiquitin thioesterase 14) (Ubiquitin-specific-processing protease 14) | USP14 TGT | C.TESEEEEVTK#GKENQLQLSCFINQEVK.Y |
| P04004 | Vitronectin (VN) (S-protein) (Serum-spreading factor) (V75) [Cleaved into: Vitronectin V65 subunit; Vitronectin V10 subunit; Somatomedin-B] | VTN | C.TEGFNVDKK#CQCDELCSYYQSCCTDYTAECKPQVTR.G |
| P04004 | Vitronectin (VN) (S-protein) (Serum-spreading factor) (V75) [Cleaved into: Vitronectin V65 subunit; Vitronectin V10 subunit; Somatomedin-B] | VTN | R.CTEGFNVDKK#C*QC*DELC*SYYQSCCTDYTAECKPQVTR.G |
| Q15942 | Zyxin (Zyxin-2) | ZYX | C.EDCGK#PLSIEADDNGCFPLDGHVLCR.K |

End of table

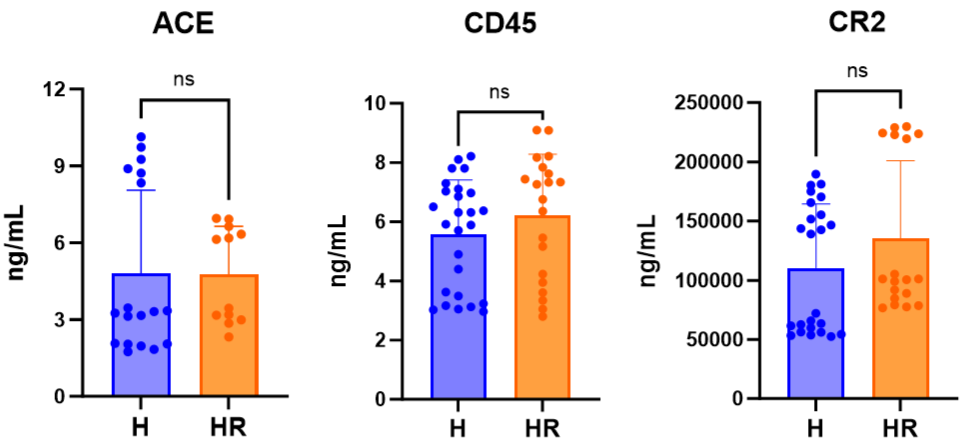


**Figure S1:** ELISA measurements of 3 of the 12 selected candidate biomarkers that did not show a change between the high-risk subjects (HR; $n$ = 3) and healthy controls (H; $n$ = 4). Data were shown in bar graphs as means ± standard deviation (SD). To ensure reproducibility of the results, measurements were conducted in replicates with various dilution factors and repeated on 2 – 5 separate occasions in aliquots of the individual samples from each subject group. Repeated measurements were overlayed on the bar graphs. Pairwise statistical comparisons were by two-tailed unpaired t-test; 'ns' indicates $p$-value > 0.05.

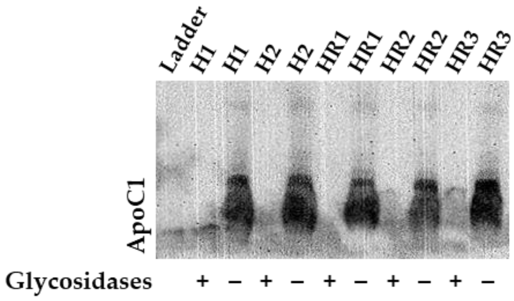


**Figure S2:** Western blot analysis of ApoC1 glycosylation. The original plasma samples from the healthy (H1 – H4) and high-risk (HR1 – HR3) subject groups in our small cohort were treated with deglycosylases (PNGase F and O-Glycosidase; New England Biolabs; Ipswich, MA, USA) prior to Western blotting. While there was a faint band of ApoC1 in one of the treated samples (H1), there was complete loss of the ApoC1 band in the remaining treated samples for reasons we could not resolve despite several attempts at troubleshooting (see the discussion for more details). The samples were treated as follows: using proprietary buffers (New England Biolabs; Ipswich, MA, USA), plasma were denatured at 100 $^{o}$C for 10 min. After this initial denaturation step, GlycoBuffer 2 and NP-40 (New England Biolabs; Ipswich, MA, USA) were added to the denatured samples, which were then divided into two equal portions; one received the deglycosylases according to the manufacturer instructions and the other portion did not. Both portions were then incubated at 37 $^{o}$C for 4 h. Shortly thereafter, the treated and untreated samples were loaded in parallel onto separate lanes/wells of a 12% Bis-Tris PAGE gel (GenScript; Piscataway, NJ, USA) and run as described in the methods section.

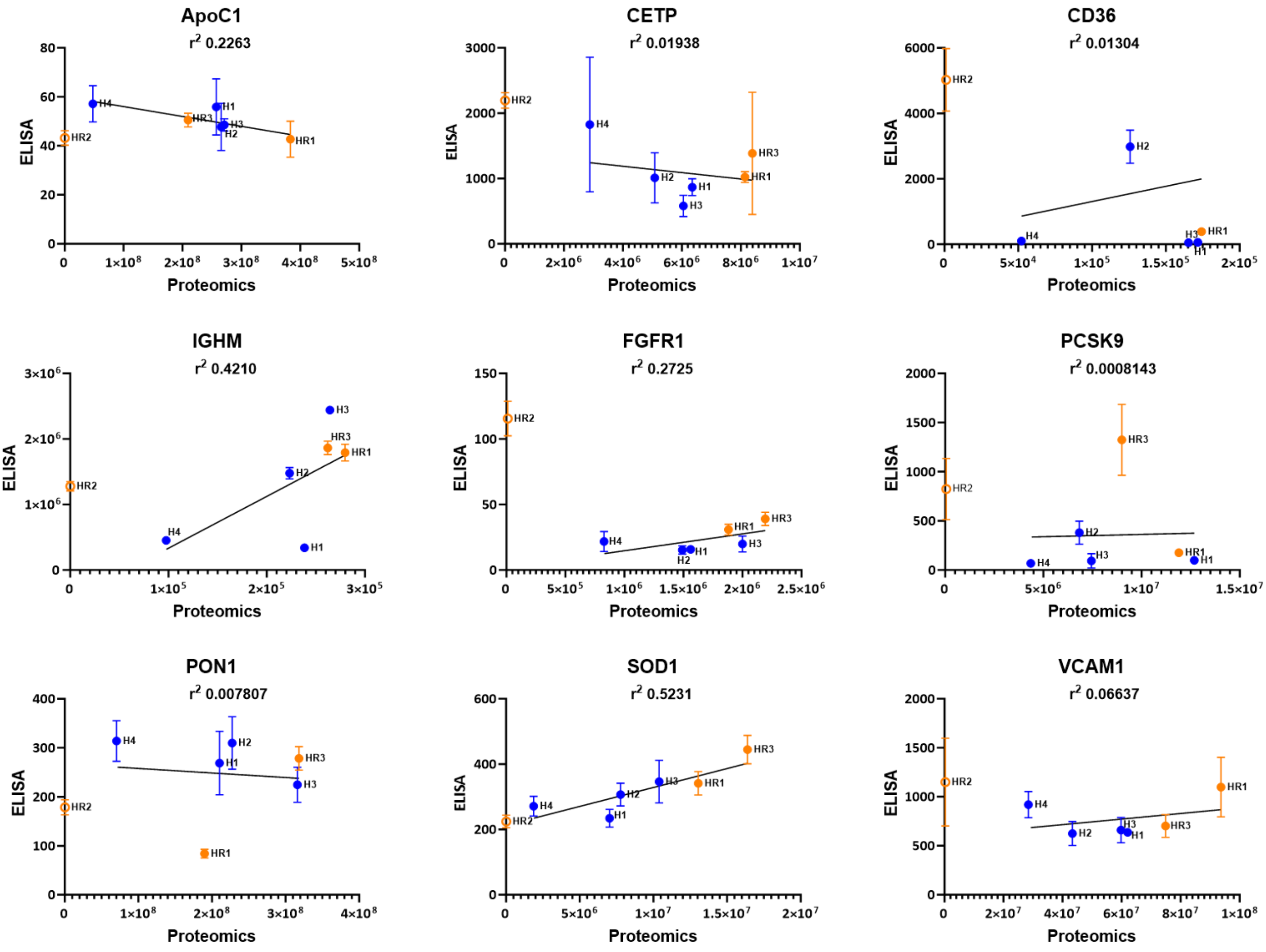


**Figure S3:** Correlation analysis of the ELISA and LC-MS/MS proteomics measurements of the 9 biomarkers that showed significant changes in the high-risk subjects compared to healthy controls among the selected 12 candidates as was detailed in the results. Simple linear regression analysis was applied to the data points (solid black lines) with goodness-of-fit ($r^2$) values shown for each. Both ELISA and LC-MS/MS measurements were performed in the original plasma samples from the subject groups in our small cohort of healthy (H1 – H4) and high-risk (HR1 – HR3) subjects, which were also utilized in our prior quadra-omics studies. For purposes of this correlation analysis, one of the high-risk subjects (HR2) was left out of the analysis as an outlier. This resulted in a positive correlation between the ELISA and LC-MS/MS measurements of 6 out of the 9 biomarkers (see the discussion for more details).
